# FMO rewires metabolism to promote longevity through tryptophan and one carbon metabolism

**DOI:** 10.1101/2021.06.18.449022

**Authors:** Hyo Sub Choi, Ajay Bhat, Marshall B. Howington, Megan L. Schaller, Rebecca Cox, Shijiao Huang, Safa Beydoun, Hillary A. Miller, Angela M. Tuckowski, Joy Mecano, Elizabeth S. Dean, Lindy Jensen, Daniel A. Beard, Charles R. Evans, Scott F. Leiser

## Abstract

Flavin containing monooxygenases (FMOs) are promiscuous enzymes known for metabolizing a wide range of exogenous compounds. In *C. elegans*, *fmo-2* expression increases lifespan and healthspan downstream of multiple longevity-promoting pathways through an unknown mechanism. Here, we report that, contrary to its classification as a xenobiotic enzyme, *fmo-2* expression leads to rewiring of endogenous metabolism principally through changes in one carbon metabolism (OCM). Using computer modeling, we identify decreased methylation as the major OCM flux modified by FMO-2 that is sufficient to recapitulate its longevity benefits. We further find that tryptophan is decreased in multiple mammalian FMO overexpression models and is a validated substrate for FMO enzymes. Our resulting model connects a single enzyme to two previously unconnected key metabolic pathways and provides a framework for the metabolic interconnectivity of longevity-promoting pathways such as dietary restriction. FMOs are well-conserved enzymes that are also induced by lifespan-extending interventions in mice, supporting a conserved and critical role in promoting health and longevity through metabolic remodeling.

## Introduction

Flavin-containing monooxygenases (FMOs) are a family of enzymes that oxygenate substrates with nucleophilic centers, such as nitrogen and sulfur^1^. They were first discovered 50 years ago and have been studied extensively under the context of xenobiotic and drug metabolism^1^. FMOs bind to an FAD prosthetic group and interact with an NADPH cofactor to oxygenate substrates^2^. The FMO protein family is highly conserved both genetically and structurally from bacteria to humans^2,3^. Considering the conserved nature of FMOs, it is plausible that they share an endogenous, more ancient physiological role than detoxifying xenobiotics.

Through a screen of genes downstream the hypoxia-inducible factor-1 (HIF-1), a longevity-promoting transcription factor in *C. elegans*, flavin-containing monooxygenase-2 (*fmo-2*) was identified as necessary for the longevity and health benefits of both hypoxia and dietary restriction (DR)^4^. The *fmo-2* gene is also sufficient to confer these benefits on its own when overexpressed^4^. Recently, studies also suggest potential endogenous role(s) for mammalian FMOs, where changes in expression of multiple FMO proteins affect systemic metabolism^5–10^. Initial correlative reports also link FMOs to the aging process, showing that *Fmo* genes are frequently induced in long-lived mouse models, such as DR mice^5,6^. However, the mechanism(s) for how *Fmos* modulate endogenous metabolism and/or aging *in vivo* is unclear, as is their potential to benefit health and longevity in multiple species.

While frequently implicated in cancer cells, recent studies identify one carbon metabolism (OCM) as a common downstream target of multiple longevity pathways^11–14^. OCM is an important intermediate metabolic pathway and refers to a two-cycle metabolic network including the folate cycle and the methionine cycle^15^. OCM takes nutrient inputs, including glucose and vitamin B12, and utilizes them to synthesize intermediates for metabolic processes involved in growth and survival, including nucleotide metabolism, the transsulfuration and transmethylation pathways, and lipid metabolism^12,13,16^. In particular, suppressing expression of the methionine cycle gene *sams-1* by RNA-mediated interference (RNAi) extends the wild type worm lifespan, but fails to further extend the lifespan of the genetic DR model *eat-2* mutants^17^.

Kynurenine synthesis from tryptophan and subsequent metabolism is another important metabolic pathway that can play a role in many processes, including longevity regulation. Knocking out tryptophan 2,3-dioxygenase (TDO), which catalyzes the first and rate-limiting step of this pathway, leads to lifespan extension in worms and flies^18,19^. Similarly, suppressing the kynurenine pathway by knocking down kynureninase (*kynu-1*) in worms also increases lifespan^20^. The kynurenine pathway competes for tryptophan with the serotonergic biosynthesis pathway and produces nicotinamide adenine dinucleotide (NAD) and other metabolites, including kynurenic acid and picolinic acid^21^.

Given that 1) induction of Fmos correlates with increased longevity across species, 2) nematode *fmo-2* is necessary and sufficient to improve health and longevity downstream of metabolic perturbations, and 3) loss of Fmo expression can modify aspects of metabolism, we hypothesized that Fmos affect aging by modifying one or more distinct metabolic processes. Therefore, we sought to determine the metabolic changes that occur when the expression of nematode *fmo-2* is perturbed to identify its mechanism of longevity regulation. Our resulting data support a model where *fmo-2* oxygenates tryptophan, leading to alteration of OCM components that confer longevity and healthspan benefits by reducing flux through methylation processes.

## Results

### Fmo-2 alters one carbon metabolism

Based on the conserved enzymatic mechanism^2,3^ and our published data supporting a key role for nematode FMO-2 in regulating stress resistance, healthspan and longevity^4^, we hypothesized that FMO-2 may significantly alter endogenous metabolism in *C. elegans*. To test if systemic metabolism was broadly altered by FMO-2, we used untargeted metabolomics analysis (**Supplementary Data 1**) of three strains with varying *fmo-2* expression: the wild type reference strain (N2 Bristol) the *fmo-2(ok2147)* putative knockout strain (FMO-2 KO), and our previously published long-lived *fmo-2* overexpression (KAE9) strain (FMO-2 OE). The resulting principal component analysis (PCA) shows a substantial explained variance (65.3%) through principal components (PC) 1 and 2 (**Figure 1A**). Our untargeted metabolomics data suggest a distinct difference in the metabolome between the three strains, consistent with expression of nematode *fmo-2* being sufficient to modify endogenous metabolism (**Figure 1B**).

**Figure 1:**
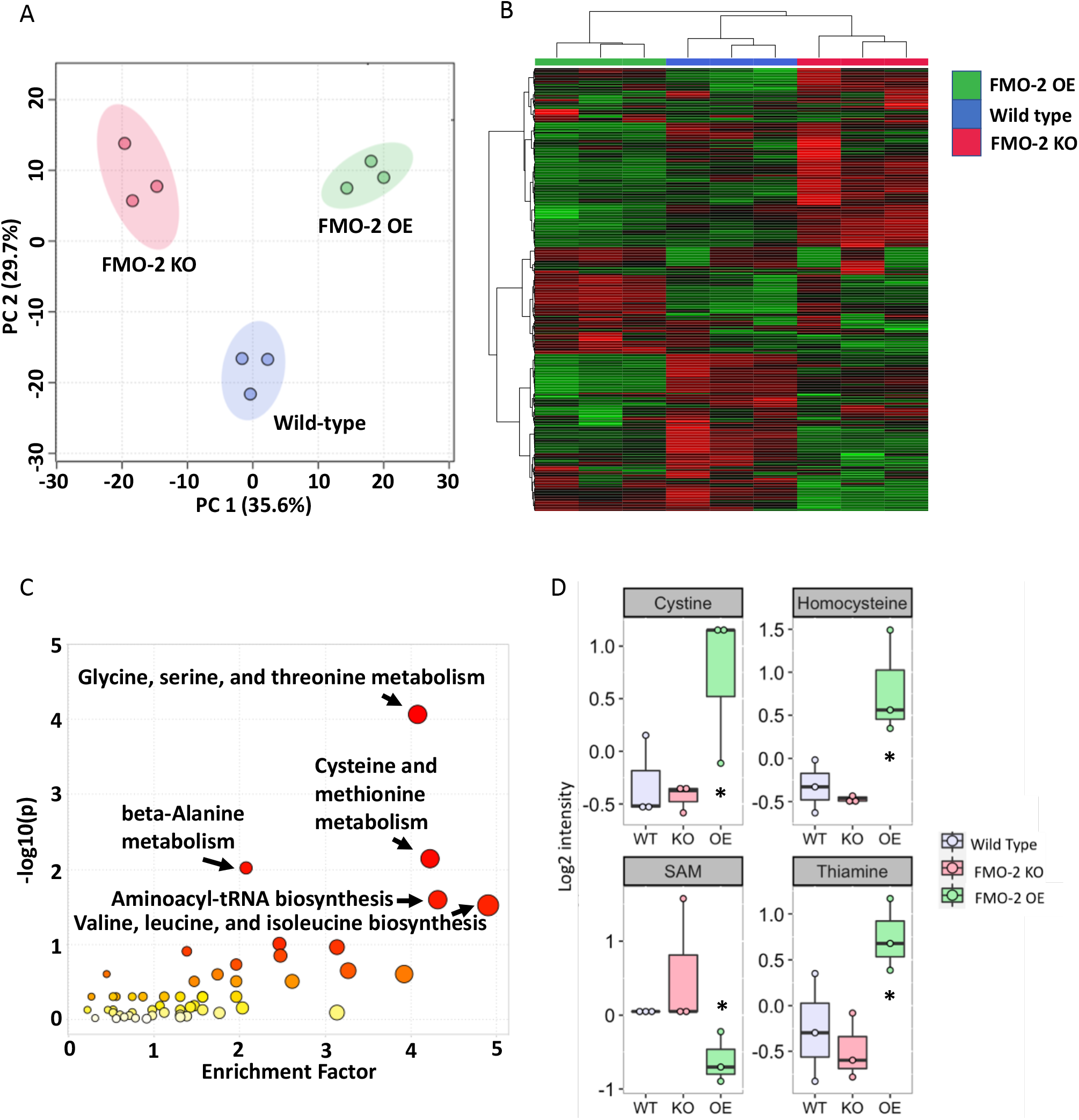
One carbon metabolism is altered by *fmo-2* expression level. A) Principal component analysis of untargeted LC-MS metabolomics data of wild type, FMO-2 OE, and FMO-2 KO strains of *C. elegans*. B) Heatmap of untargeted LC-MS metabolomics data of the wild type, FMO-2 OE and FMO-2 KO. C) Pathway enrichment analysis using untargeted LC-MS metabolomics data of wild type and FMO-2 OE. D) Comparison of targeted metabolomics data of metabolites related to OCM between the wild type, FMO-2 OE and FMO-2 KO normalized to the median and log transformed. SAM = s-adenosylmethionine. * represents p < 0.05 using paired t-test.

Having established broadly that *fmo-2* expression modifies metabolism, we next asked what key metabolic aspects are modified. Using p-value < 0.05 as our significance threshold, we identified five metabolic pathways that are significantly altered by the overexpression of *fmo-2*, most of which are involved in amino acid metabolism (**Figure 1C, Supplementary Data 2**). Of the five pathways, we observed the most significant enrichment in glycine, serine, and threonine metabolism (**Figure 1C**). Exogenous supplementation of glycine in worm diet is reported to extend lifespan by remodeling the methionine cycle^22^, a component of one carbon metabolism (OCM) and another significantly enriched metabolic pathway from our analysis, cysteine and methionine metabolism (**Figure 1C, Supplementary Data 2**). Indeed, OCM is a nexus of multiple metabolic pathways that are necessary for survival; OCM is implicated in multiple longevity pathways, including dietary restriction, insulin/IGF-1 signaling, and the metformin-induced longevity response^13,16,23^. Due to its relevance in multiple longevity pathways and the direct involvement of cysteine and methionine metabolism within this metabolic network, we postulated that *fmo-2* regulates longevity through its interactions with OCM.

To test whether *fmo-2* expression modifies OCM, we used targeted metabolomics analysis on a panel of metabolites involved in OCM and related pathways to determine whether their abundance levels were altered following *fmo-2* expression (**Supplementary Data 3**). We hypothesized that the affected metabolites would have abundance levels that correlate with *fmo-2* expression level. Thus, we compared the level of metabolite abundance between the wild type and FMO-2 OE and also between the wild type and FMO-2 KO. Consistent with our hypothesis that OCM is altered by *fmo-2* expression, we observed significant changes in the abundance level of cystine, homocysteine, s-adenosylmethionine (SAM), and thiamine in FMO-2 OE compared to the wild type (**Figure 1D**). With the exception of cystine, we observed insignificant but consistent trends in the abundance level of these metabolites that moved in the opposite direction in FMO-2 KO compared to FMO-2 OE (**Figure 1D**). The insignificant trends in FMO-2 KO are consistent with our previous observation that knocking out *fmo-2* does not affect worm lifespan^4^. Similarly, we also observed insignificant but consistent trends in the abudance level that correlate with *fmo-2* expression level in multiple other metabolites that we measured (**Supplementary Figure 1**). Taken together, our results are consistent with the hypothesis that the OCM pathway is modified by *fmo-2* expression.

**Supplementary Figure 1:**
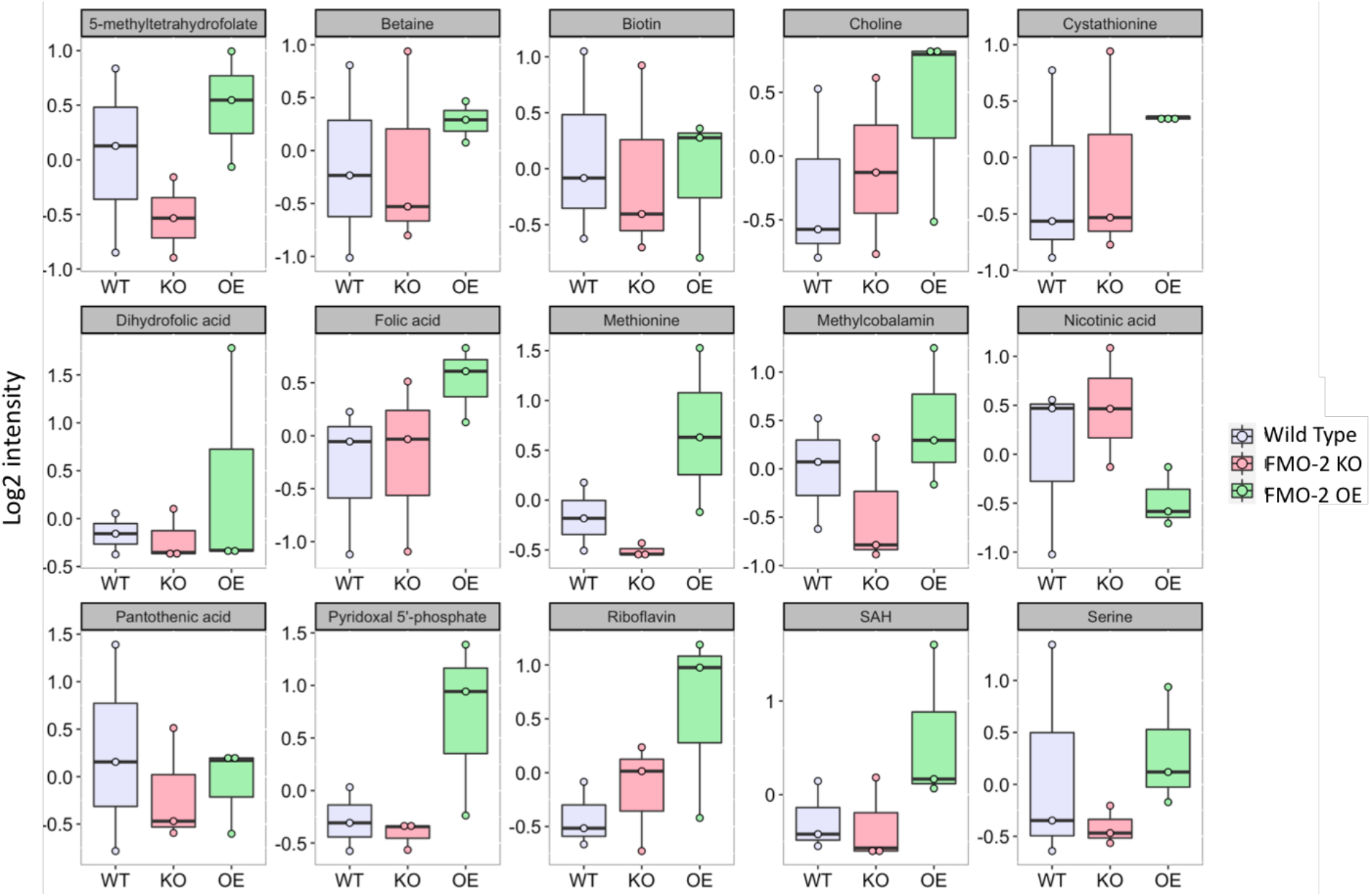
Comparison of targeted metabolomics data of metabolites related to OCM between the wild-type, FMO-2 OE and FMO-2 KO. SAH = s-adenosylhomocysteine. Data are median normalized and log transformed.

### One carbon metabolism interacts with fmo-2 to regulate stress resistance and longevity

Having established that FMO-2 modifies endogenous metabolism broadly and OCM specifically, we next hypothesized that these metabolic changes are causal for longevity phenotypes. Previous studies identify increased stress resistance as a common phenotype shared by multiple long-lived organisms both within and between species^24–27^. To determine the functional interaction between *fmo-2* and OCM, we used RNAi to knockdown the expression of genes involved in OCM (**Figure 2A**) and tested for their role in promoting or repressing survival against the oxidative stressor paraquat. Individual knockdown of multiple genes exhibit altered paraquat stress resistance phenotypes for the wild type and FMO-2 OE. Of the eight genes that we tested, the knockdown of five genes, *alh-3*, *atic-1*, *cbs-1*, *cth-2*, and *sams-1*, abrogate FMO-2 OE resistance against paraquat (**Figure 2B-F**), as assessed using log-rank test with a cutoff threshold of p < 0.0001^28^ compared to the empty vector (EV) controls. Our data for the knockdowns of *alh-3*, *atic-1*, and *cbs-1* are consistent with previous reports that their expression levels are upregulated in long-lived worms^11,29^. *alh-3* is upregulated in *eat-2* mutants, *atic-1* is upregulated in both *eat-2* and *daf-2* mutants, and *cbs-1* is upregulated under cold-induced longevity and is required for the lifespan extension of *eat-2* and *glp-1* mutants^11,29–31^. *cth-2* is a homolog of the transsulfuration pathway enzyme cystathionine *γ*-lyase that is detrimental to the wild type lifespan when its expression is suppressed using RNAi^30^. Thus, it is plausible that these genes are required for multiple longevity pathways, including *fmo-2*-mediated longevity, to confer resistance against paraquat. Interestingly, while *sams-1* knockdown extends worm lifespan^17^, we find that knocking down *sams-1* abrogates FMO-2 OE paraquat resistance (**Figure 2F**), suggesting that the regulation of lifespan and stress resistance are uncoupled in this instance. This result is similar to previous work showing that *sams-1* knockdown is detrimental to survival under pathogen exposure^32^.

**Figure 2:**
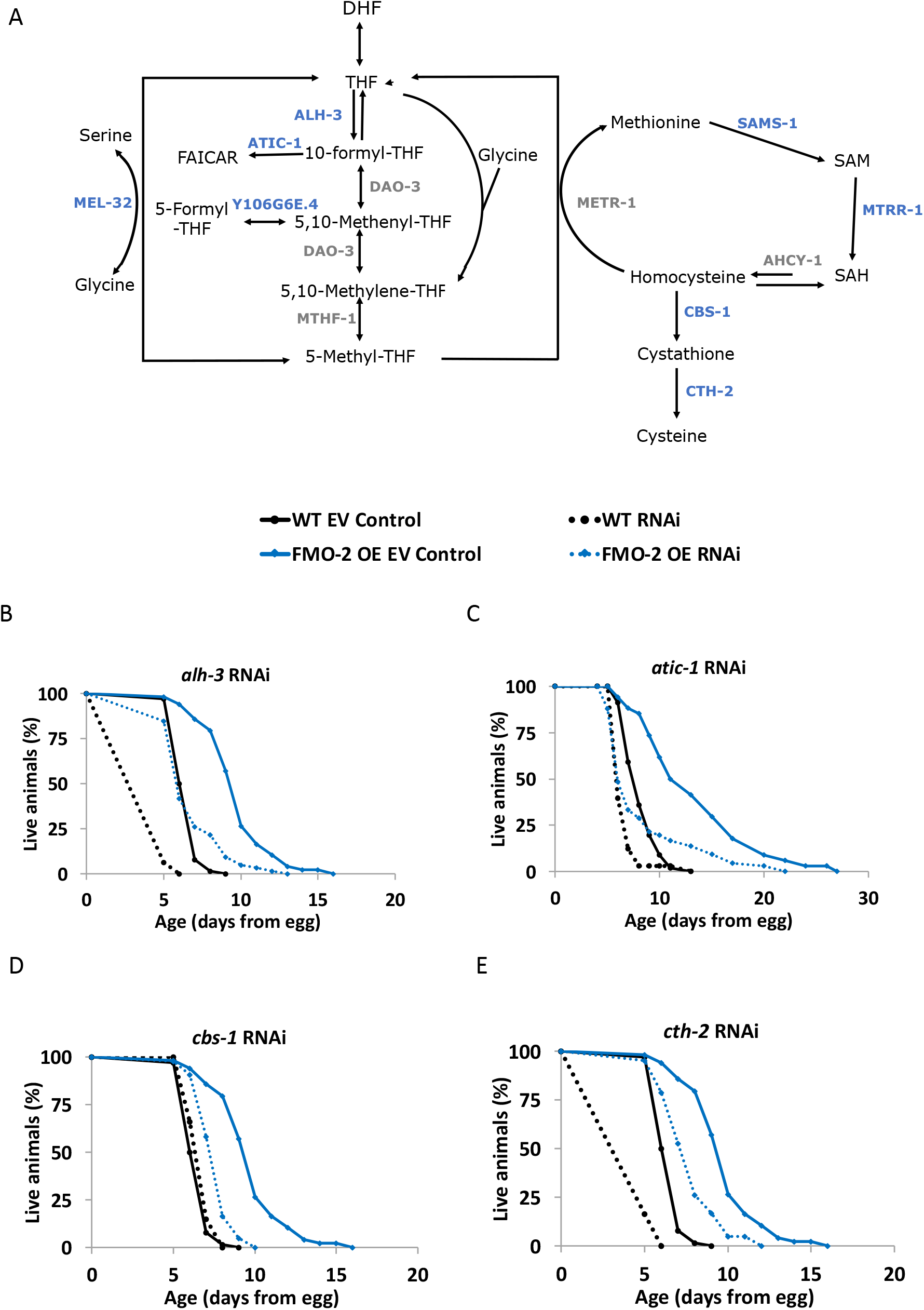
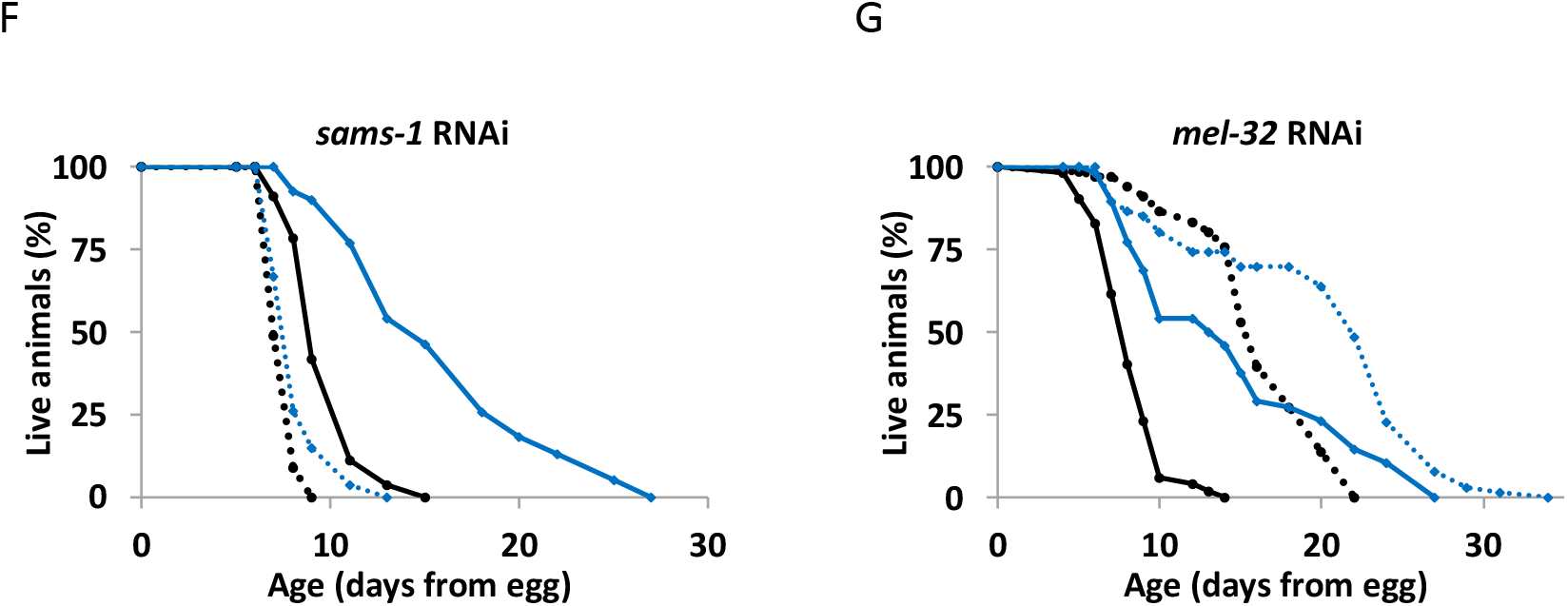
*Fmo-2* interacts with OCM genes to regulate resistance against paraquat. A) Diagram of OCM network. Genes included in the screen are labeled in blue and genes not included in the screen are labeled in gray. B-G) 5 mM paraquat stress resistance assay (from L4) comparing the wild-type and FMO-2 OE on empty-vector (EV) and B) *alh-3* RNAi, C) *atic-1* RNAi, D) *cbs-1* RNAi, E) *cth-2* RNAi, F) *sams-1* RNAi, and G) *mel-32* RNAi. Statistics in Supplementary Data 4.

We also find that knocking down *mel-32* increases stress resistance of both the wild type and FMO-2 OE (**Figure 2G**), suggesting that the stress resistance conferred by the suppression of *mel-32* is independent of *fmo-2*. The two remaining genes that we knocked down, *mtrr-1* and Y106G6E.4, did not affect the stress resistance of the worms (**Supplementary Figure 2**). Overall, our data show that five out of eight of the genes that we tested are required by FMO-2 OE to confer paraquat resistance. These results are consistent with the hypothesis that OCM is a regulator of stress resistance and that there is a genetic interaction between *fmo-2* and OCM in that regulation.

**Supplementary Figure 2:**
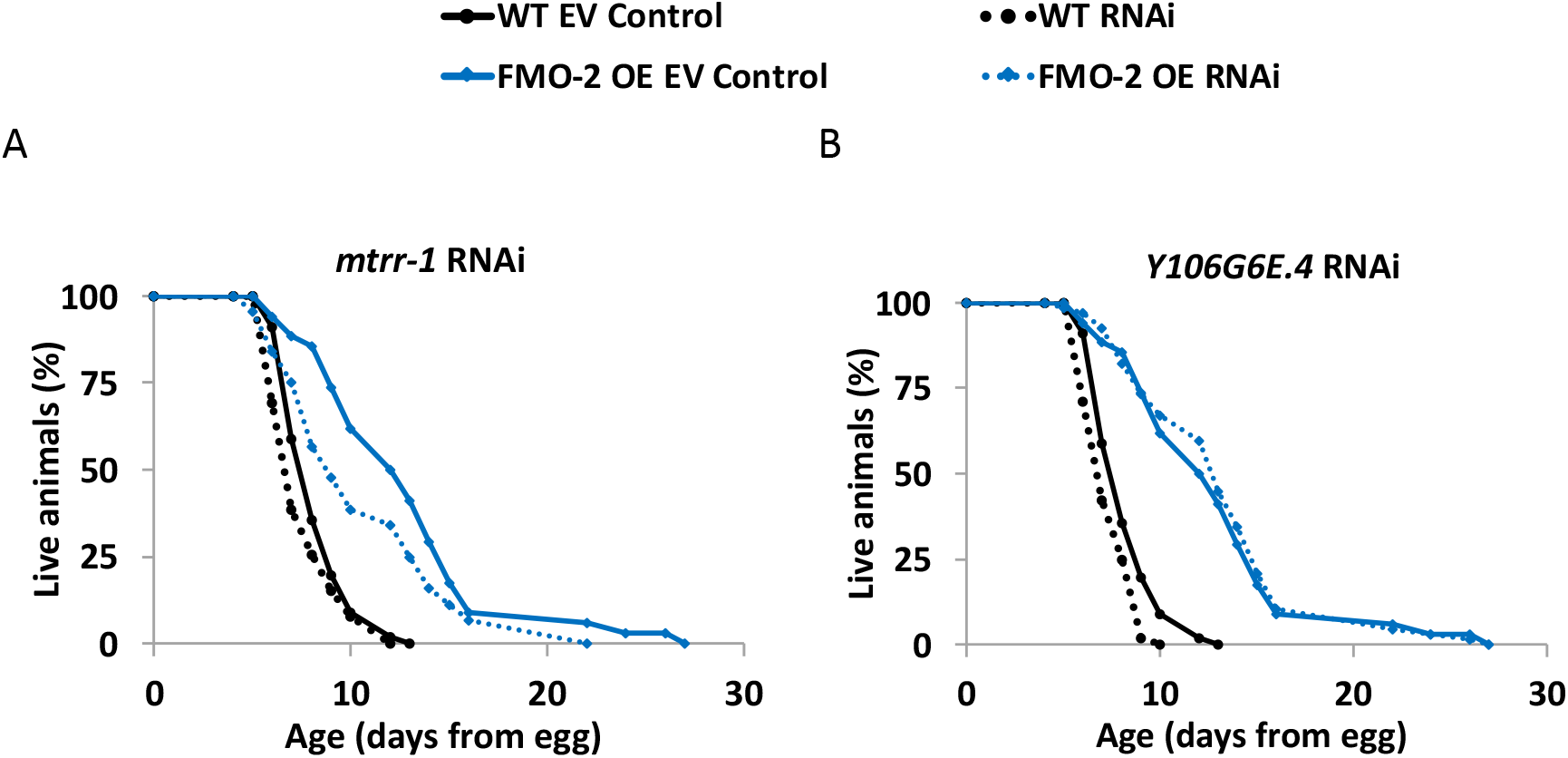
OCM genes that do not alter worm stress resistance. 5 mM paraquat stress resistance assay comparing the wild-type and FMO-2 OE on empty-vector (EV) and A) *mtrr-1* RNAi and B) *Y106G6E.4* RNAi. Statistics in Supplementary Data 4.

To test the interaction between *fmo-2* and OCM more directly, we measured the longevity of worms with RNAi knockdown of genes from our paraquat resistance screen. We included FMO-2 KO in the lifespan analysis to determine if the interactions that we identify are dependent on *fmo-2* expression. Similar to the paraquat survival assays, multiple gene knockdowns showed altered lifespan phenotypes for the wild type, FMO-2 OE, and FMO-2 KO. Of the eight genes we tested, knockdown of two genes, *alh-3* and *cth-2*, suppress the lifespan extension of FMO-2 OE to the level of the wild type (**Figure 3A, B**), as assessed using log-rank test with a cutoff threshold of p < 0.0001^28^ compared to the empty vector (EV) controls, consistent with our paraquat survival data. In contrast, knockdown of *sams-1* increases the lifespan of the wild type and FMO-2 KO to the level of FMO-2 OE without further extending the lifespan of FMO-2 OE (**Figure 3C**), suggesting a separation between lifespan and stress resistance. Additionally, we find that knocking down *mel-32* only extends the lifespan of FMO-2 OE (**Figure 3D**). It is possible that the metabolic alterations caused by increased *fmo-2* expression are required for *mel-32* gene suppression to promote worm lifespan.

**Figure 3:**
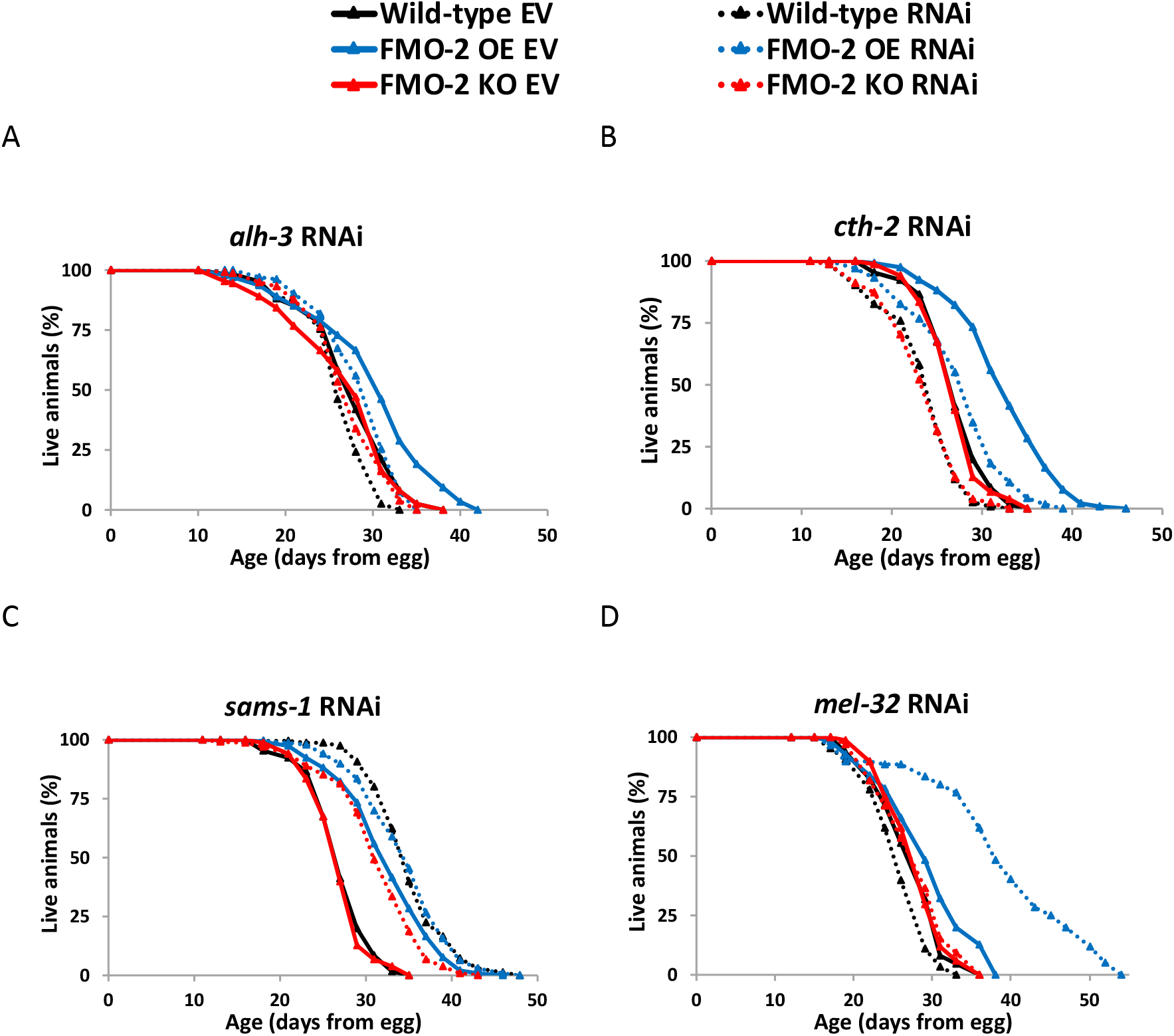
*Fmo-2* interacts with OCM genes to regulate lifespan. Lifespan analysis comparing the wild-type and FMO-2 OE on empty-vector (EV) and A) *alh-3* RNAi, B) *cth-2* RNAi, C) *sams-1* RNAi and D) *mel-32* RNAi. Statistics in Supplementary Data 5.

Knocking down the remaining four genes, *atic-1*, *cbs-1*, *mtrr-1*, and Y106G6E.4, do not affect the lifespan of any of the worm strains (**Supplementary Figure 3**). Although knocking down *atic-1* and *cbs-1* abrogated FMO-2 OE paraquat resistance (**Figure 2C, D**), they did not abrogate FMO-2 OE lifespan (**Supplementary Figure 3A, B**). Similar to *sams-1* knockdown data, this finding suggests uncoupling of stress resistance and lifespan. In total, our data show that two of the genes that we tested are required for FMO-2 OE lifespan extension, another gene extends lifespan non-additively with FMO-2 OE, placing it in the same functional pathway, and one of the genes only extends the lifespan of FMO-2 OE when knocked down. Thus, our data support an interaction between *fmo-2* and genes involved with OCM in regulating worm lifespan.

**Supplementary Figure 3:**
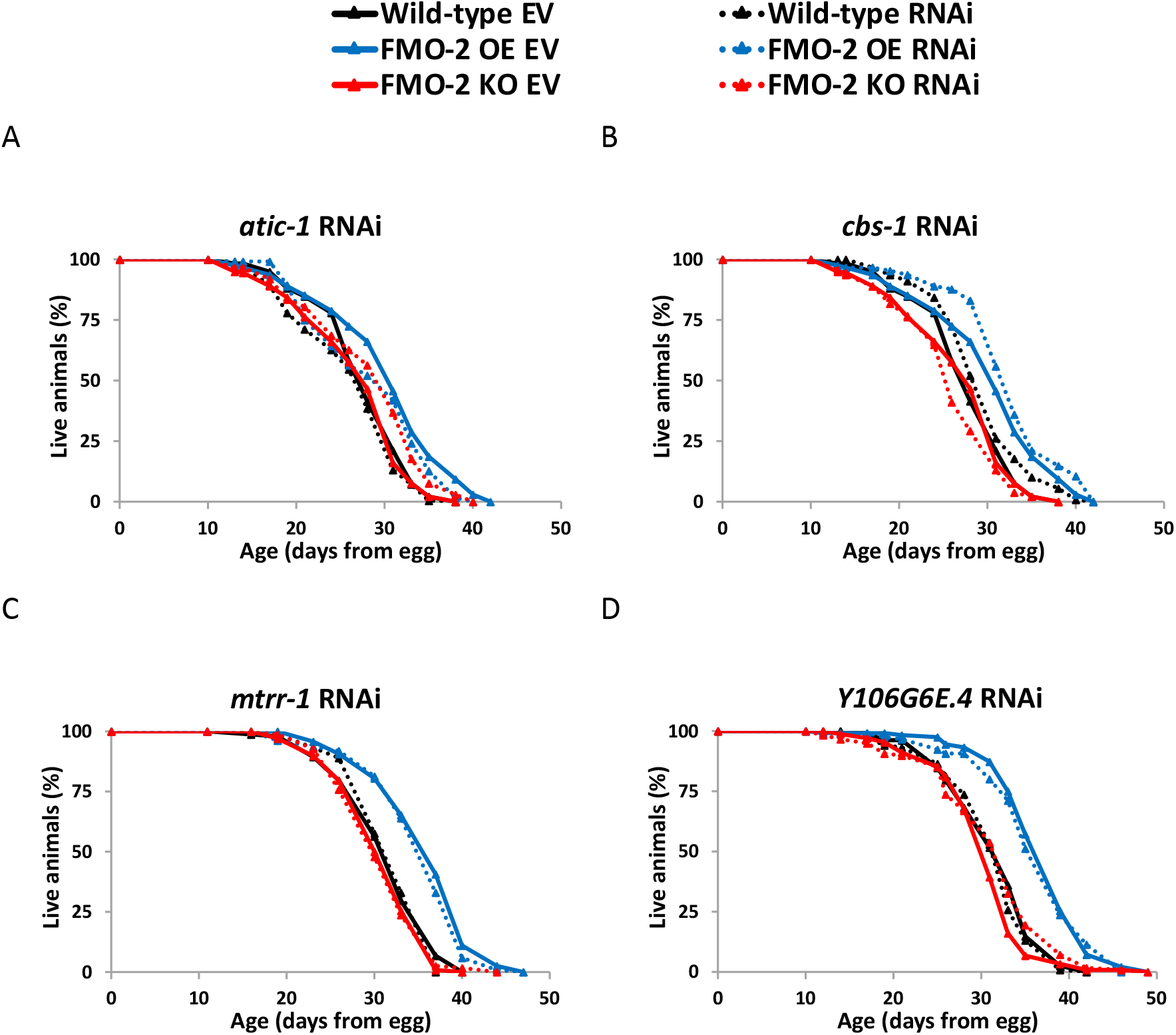
OCM genes that do not alter worm lifespan. Lifespan analysis comparing the wild-type and FMO-2 OE on empty-vector (EV) and A) *atic-1* RNAi, B) *cbs-1* RNAi, C) *mtrr-1* RNAi, and D) *Y106G6E.4* RNAi. Statistics in Supplementary Data 5.

### Fmo-2 influences longevity by modulating the transmethylation pathway

Our data are consistent with a model where *fmo-2* interacts with OCM to regulate longevity and stress resistance. Previous studies identify multiple pathways that affect longevity and are also involved in OCM, including nucleotide metabolism, the transsulfuration pathway, and the transmethylation pathway^11,16,17^. Some of these pathways are also implicated in modifying longevity downstream of dietary restriction in multiple animal models^16,17,33^, making it likely that one or more of these pathways are in the same functional pathway as *fmo-2*. However, the metabolic consequences of *fmo-2* expression on these pathways are not clear based on the changes observed in our targeted metabolomics analysis alone, as the data only show metabolic changes at a single time point and most of the metabolites within OCM are intermediate metabolites. The stress resistance and lifespan results further complicate interpretation as some genes do not affect these phenotypes and some have effects that are independent of *fmo-2*.

To help determine the biological relevance of the changes we observed in the OCM network following *fmo-2* expression, we applied a computational model (**Supplementary Data 6**) to predict how enzyme expression (**Supplementary Data 7**) changes may affect the output fluxes of OCM. The model assumes a steady-state mass balance of fluxes in the reactions illustrated in **Figure 4A**. This simple model includes eight reaction fluxes and five fluxes representing transport of methionine (met), tetrahydrofolate (thf), s-adenosylmethionine (sam), cysteine (cys), and 5,10-methylenetetrahydrofolate (5,10thf) into and out of the folate cycle and the methionine cycle. The model output fluxes represent important inputs for the nucleotide metabolism, the transsulfuration pathway, and the transmethylation pathway, each of which are reported to be important for influencing the aging process^11,16,17^ and are potential key targets for the *fmo-2*-mediated longevity response. The stoichiometric coefficients for the reaction and transport processes in this system are stored in the matrix S (**Supplementary Data 8**), where under steady-state conditions S\***J** = **0**, where **J** is the vector of fluxes^34,35^. The entries in the vector **J** and matrix S are defined in **Figure 4A**. Vectors that satisfy the mass-balance relationship S\***J** = **0** are said to belong to the nullspace of S. To predict how changes in the expression of genes for the enzymes catalyzing the reactions in this network may affect the output fluxes, we projected the gene expression data (**Supplementary Data 7**) onto the nullspace of S (details are provided in the Methods). This projection predicts an inverse correlation between *fmo-2* expression and flux through methylation reactions, where the methylation flux is predicted to be reduced in FMO-2 OE and increased in FMO-2 KO compared to wild type (**Figure 4B, Supplementary Figure 4**). This analysis does not predict correlative changes to flux through nucleotide metabolism or the transsulfuration pathway.

**Figure 4:**
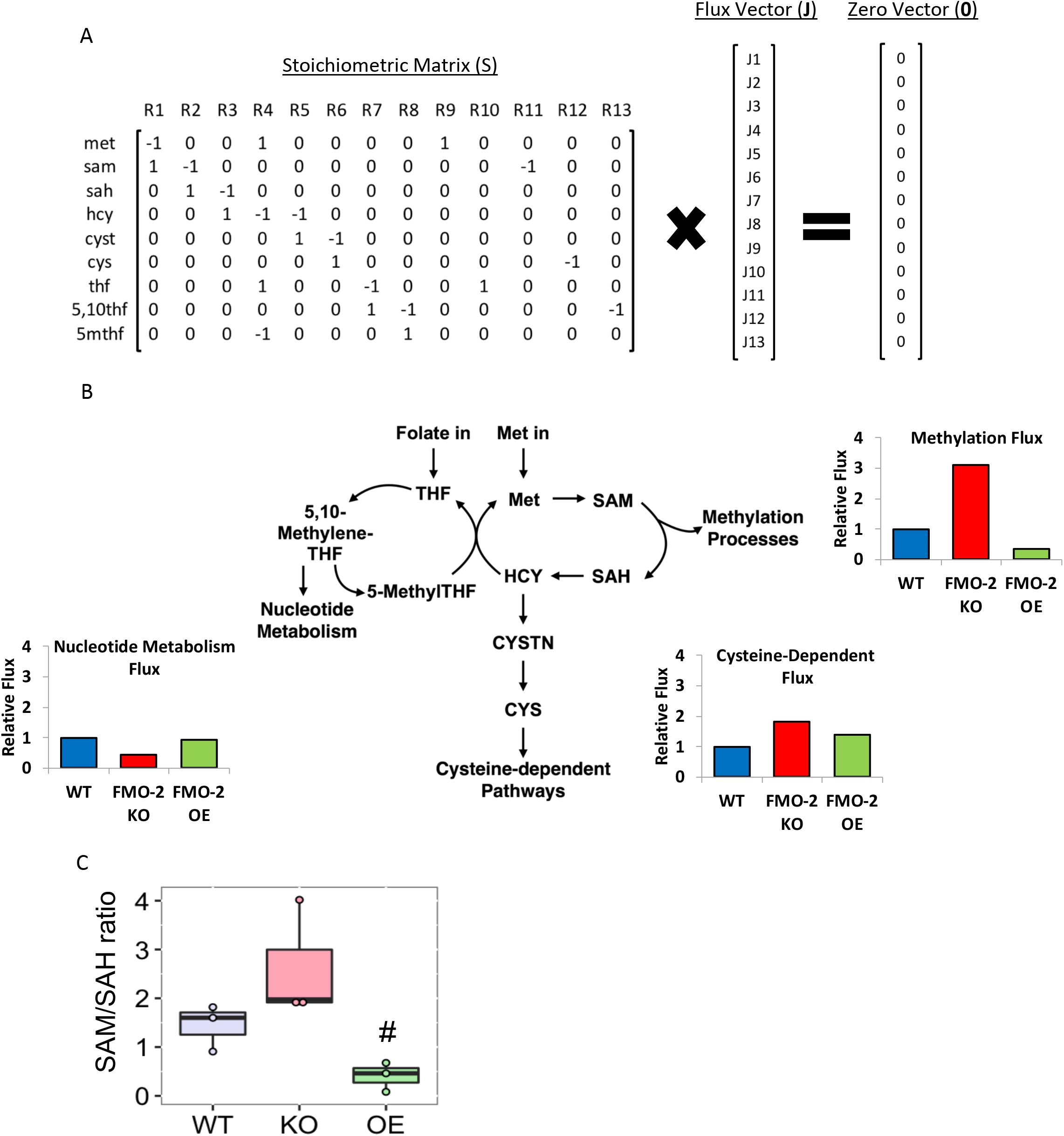
Methylation flux is altered following changes in *fmo-2* expression. A) Schematic of computational model. B) Model predictions of output metabolic fluxes. C) SAM/SAH ratio of the wild-type (WT), FMO-2 OE (OE) and FMO-2 KO (KO). # = p < 0.05 compared to the wild-type using one-way ANOVA.

Based on this analysis, we hypothesized that artificially decreasing the flux through methylation should replicate FMO-2 OE longevity in the wild type and FMO-2 KO strains, while not affecting the FMO-2 OE worms. *Sams-1* encodes for <u>s</u>-<u>a</u>denosyl<u>m</u>ethionine <u>s</u>ynthase and is involved in the conversion of methionine into s-adenosylmethionine (SAM). Suppression of *sams-*1 has been previously shown to decrease SAM level^36^ and increase longevity^17^. We find that *sams-1* RNAi recapitulates FMO-2 OE lifespan extension in the wild type and FMO-2 KO worms while not affecting FMO-2 OE lifespan **(Figure 3C)**. Our data are consistent with previous studies showing that knockdown of *sams-1* fails to further extend the lifespan of genetic DR model *eat-2* mutants^17^, thereby placing *sams-1* knockdown in the same functional pathway as FMO-2 OE.

To validate the model metabolically, we used the abundance level of SAM and s-adenosylhomocysteine (SAH) from our targeted metabolomics analysis to calculate the SAM/SAH ratio. The SAM/SAH ratio is used as a biomarker for methylation potential, where a decrease in the ratio suggests a hypomethylated state and an increase suggests a hypermethylated state^37,38^. Consistent with our computational model prediction, we observed a reduction in the SAM/SAH ratio for FMO-2 OE (hypomethylation) and an increase in the ratio for FMO-2 KO (hypermethylation) compared to the wild type (**Figure 4C**). Overall, our computational model prediction and experimental data support the hypothesis that *fmo-2* expression reduces flux through the transmethylation pathway, and that this reduction extends worm lifespan.

**Supplementary Figure 4:**
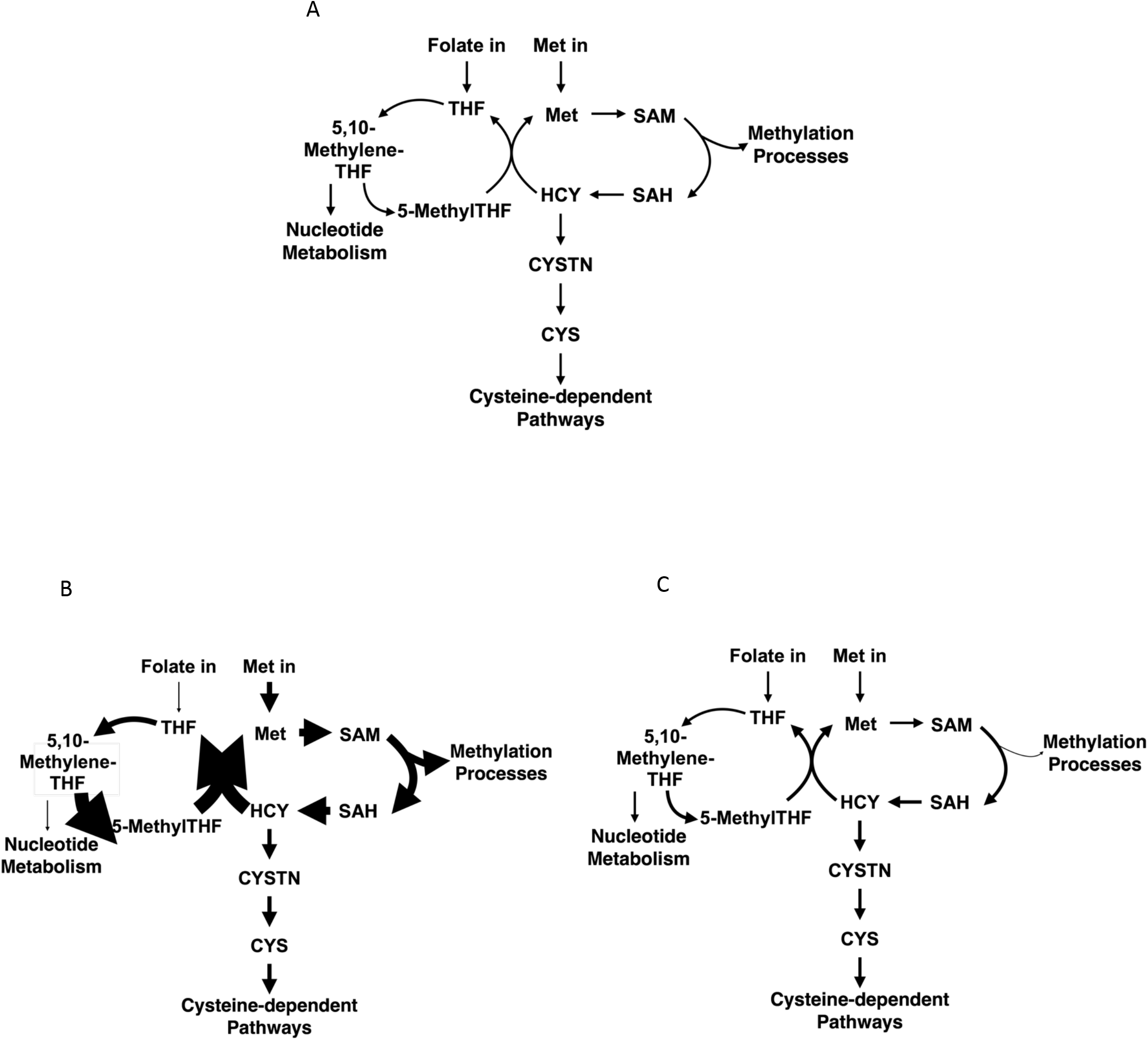
Computational model predicts reduced flux through methylation processes. Model predictions of OCM fluxes for A) the wild-type, B) FMO-2 OE, and C) FMO-2 KO after normalization to the wild-type. Arrow weights represent changes in each flux relative to the wild type, which is set to be equal to 1.

### Mammalian FMO metabolomics analysis reveals tryptophan as a substrate of FMO-2

Our data thus far suggest a model where *fmo-2* interacts with OCM to modulate the aging process. However, since FMOs are promiscuous enzymes that oxygenate many nucleophilic atoms, the mechanism by which *fmo-2* induction leads to changes in OCM is not readily evident. FMOs are known as xenobiotic metabolizing enzymes, with many known exogenous targets and few known endogenous targets^1^. Despite extensive knowledge on their enzymatic activity and recent data linking FMOs to endogenous metabolism, no link between specific and systemic metabolism has been made. We hypothesize that a limited number of FMO targets are causal in FMO-2’s effects on OCM and, importantly, on the aging process.

Due to the high degree of conservation of catalytic residues between mouse FMOs and CeFMO-2 (**Figure 5A**), we referred to our previously published targeted metabolomics of mouse FMO overexpressing (OE) HepG2 cells to determine potential metabolic targets of FMO-2^10^. Our selection criteria for putative substrates of FMO-2 included identifying metabolites that had decreased abundance in at least three of the five FMO OE cell lines to pDEST controls. We use this stringent criteria to identify the most well-conserved targets of FMOs, given that no data exist for CeFMO-2 targets. Using this approach, we identified tryptophan and phenylalanine as potential substrates of FMOs (**Figure 5B**). To determine if either of these are substrates of FMO-2, we measured the enzymatic activity of isolated FMO-2 protein in the presence of varying concentrations of tryptophan and phenylalanine. We find that FMO-2 is active toward tryptophan at a reasonable K_m_ and k_cat_ (K_m_: 880 ± 430 μM; k_cat_: 9.7 ± 1.5 sec^-1^), suggesting that tryptophan is a viable substrate of FMO-2 (**Figure 5C, Supplementary Data 9**). FMO-2 was also active toward phenylalanine, but enzymatic activity did not become apparent until 10 mM, suggesting that phenylalanine is not likely a good endogenous substrate of FMO-2 (**Figure 5D**). Since FMO-2 has no previously reported activity toward tryptophan, we used LC-MS with 100, 250, and 500 μM tryptophan under the same enzymatic conditions to determine the product of tryptophan oxygenation. Our resulting data show the presence of N-formylkynurenine in a concentration dependent manner in each of the samples, suggesting that is the product formed by FMO-2 activity toward tryptophan (**Figure 5E**). To determine whether tryptophan is a conserved substrate of FMOs, we next tested whether mFMO5 can also oxygenate tryptophan. mFMO5 also shows increased levels of N-formylkynurenine based on HPLC analysis under tryptophan enzymatic conditions (**Figure 5E**, **Supplementary Data 10**). Alignment of mFMO5 and CeFMO-2 using ancient mammalian FMO5^39–41^ shows that all but one of the catalytic residues of CeFMO-2 are conserved in mFMO5 (**Figure 5A**), so it follows that they would have similar activity toward some substrates. The kinetic parameters of FMO-2 toward NADPH, methimazole, and tryptophan are summarized in **Figure 5F**. The poor substrates (such as cysteine, phenylalanine, and TMA) and non-substrates of FMO-2 (such as 2-heptanone) are summarized with either the concentration of substrate at which FMO-2 activity is first detected or labeled not determined (N.D.) in **Supplementary Data 9**.

**Figure 5:**
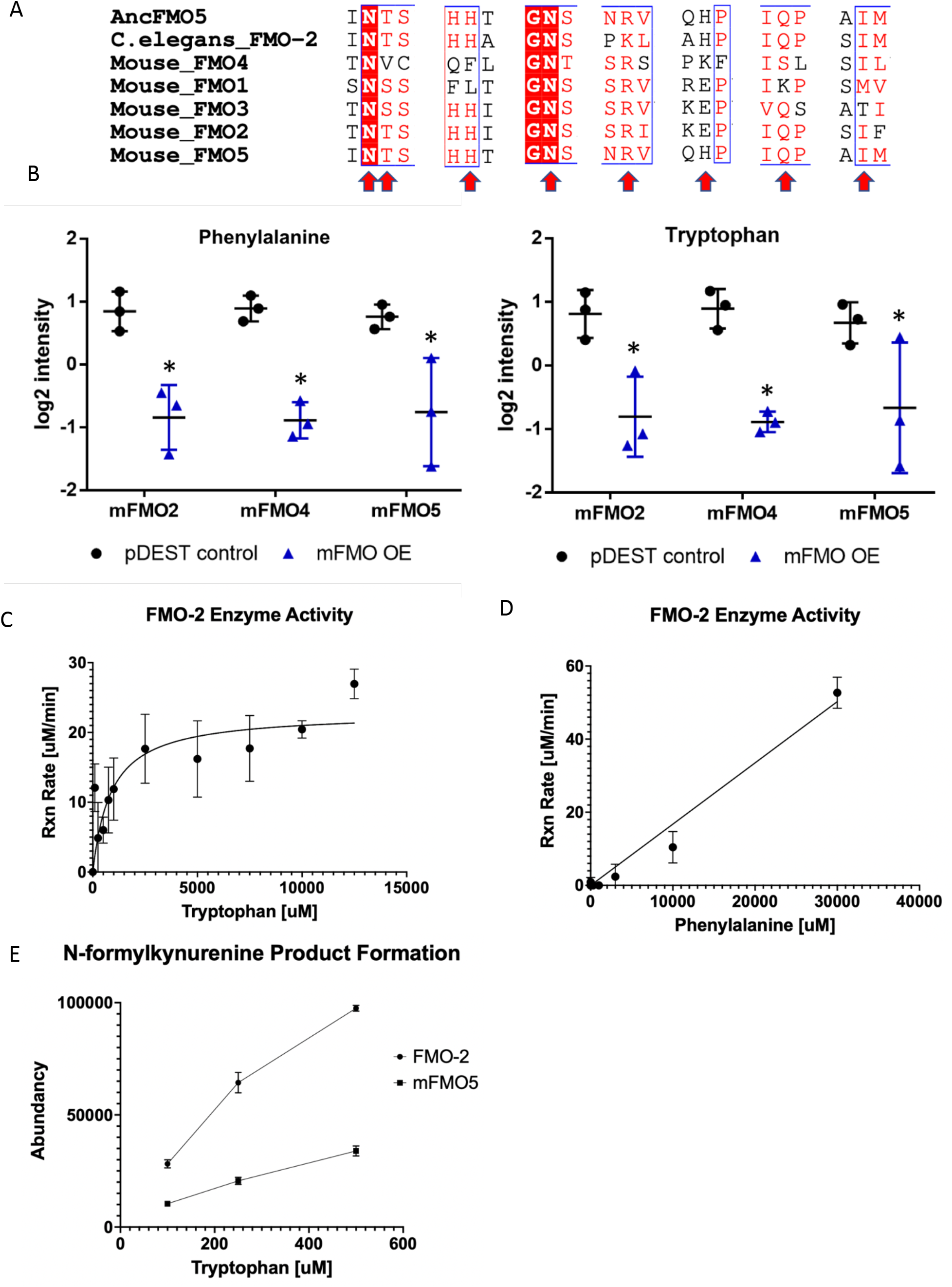
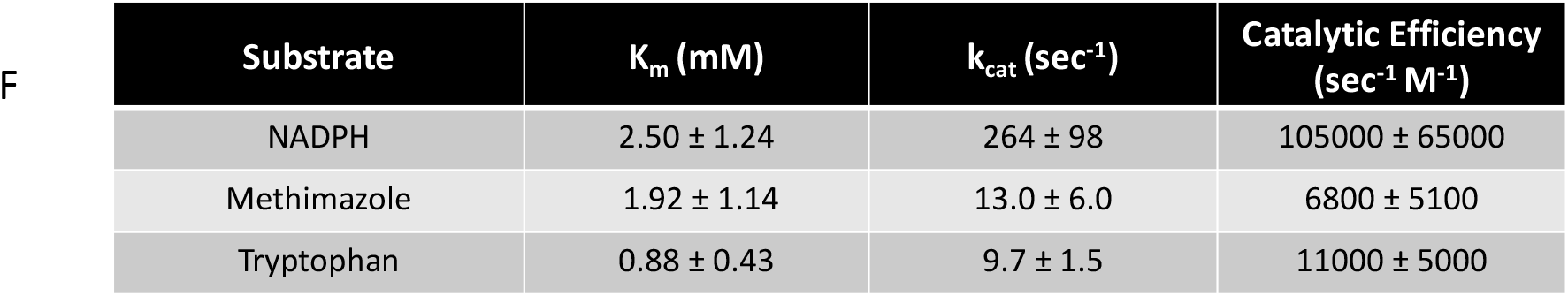
Mammalian FMO metabolomics analysis reveals the tryptophan/kynurenine pathway as a target of FMO-2. A) Conserved catalytic residues between CeFMO-2 and mouse FMOs (indicated by red arrows). B) The level of phenylalanine and tryptophan present in HepG2 cells expressing pDEST control vector, mFMO2, mFMO4, and mFMO5. * represents p < 0.05 by paired t-test. C-D) The reaction rate by concentration for purified CeFMO-2 enzyme toward tryptophan and phenylalanine at 30°C. E) The abundancy of N-formylkynurenine based on HPLC analysis of CeFMO-2 and mFMO5 activity toward 100, 250, and 500 uM tryptophan at 30°C. F) Summary table of Michaelis-Menton parameters for CeFMO-2 cofactor and substrates.

Based on our initial data linking FMO-2 to OCM, it is important to note that in addition to being a key process in the kynurenine pathway, the conversion of tryptophan to N-formylkynurenine precedes the conversion of N-formylkynurenine to kynurenine by formamidase, a process that releases formate, which is also a carbon source for OCM^42^. Formate can enter OCM through the folate cycle, thus providing a connection between tryptophan metabolism, the kynurenine pathway, and OCM. Based on this information, we hypothesize that the kynurenine pathway is a target of FMO-2 that leads to changes in OCM.

**Supplementary Figure 5:**
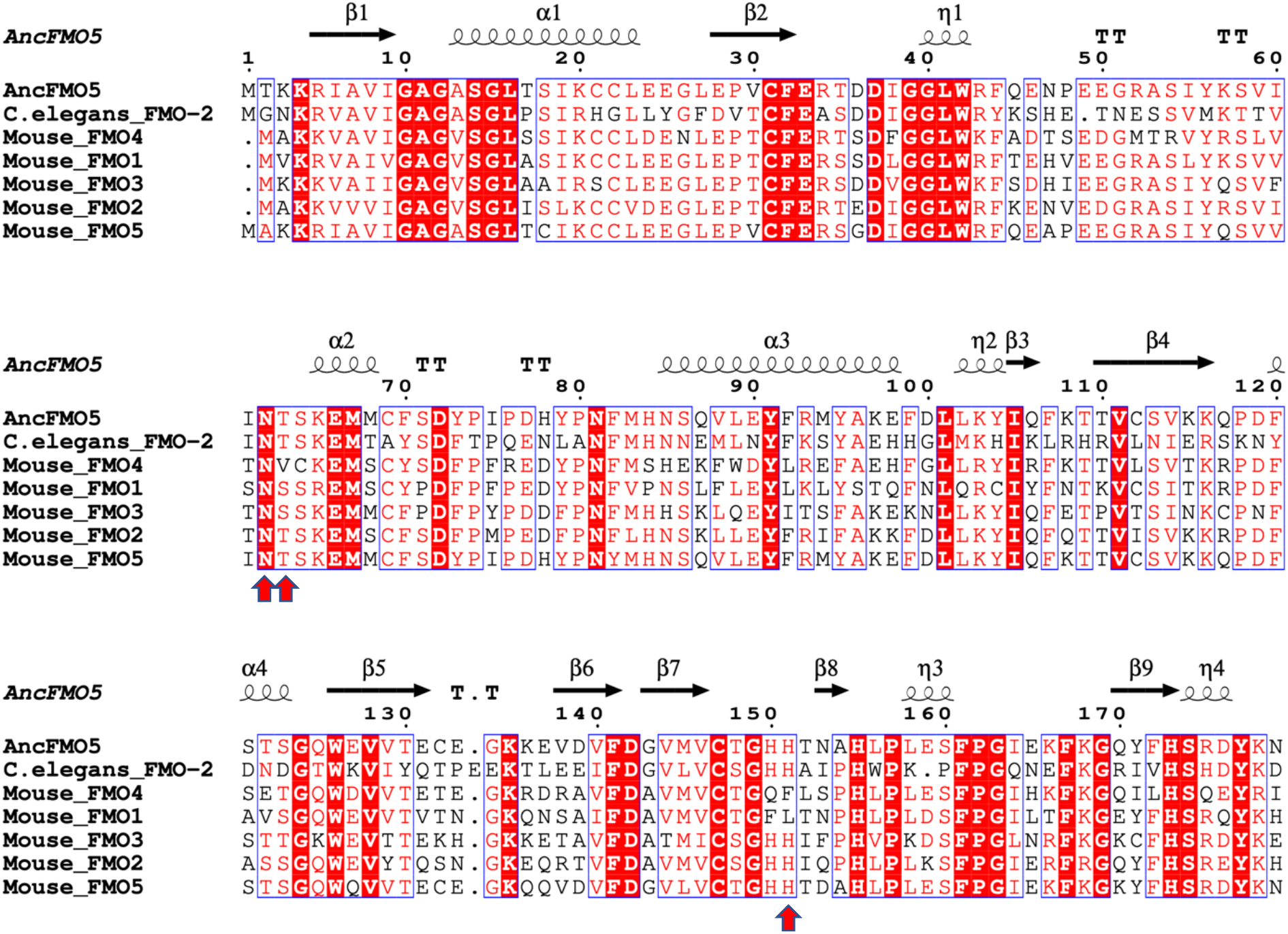
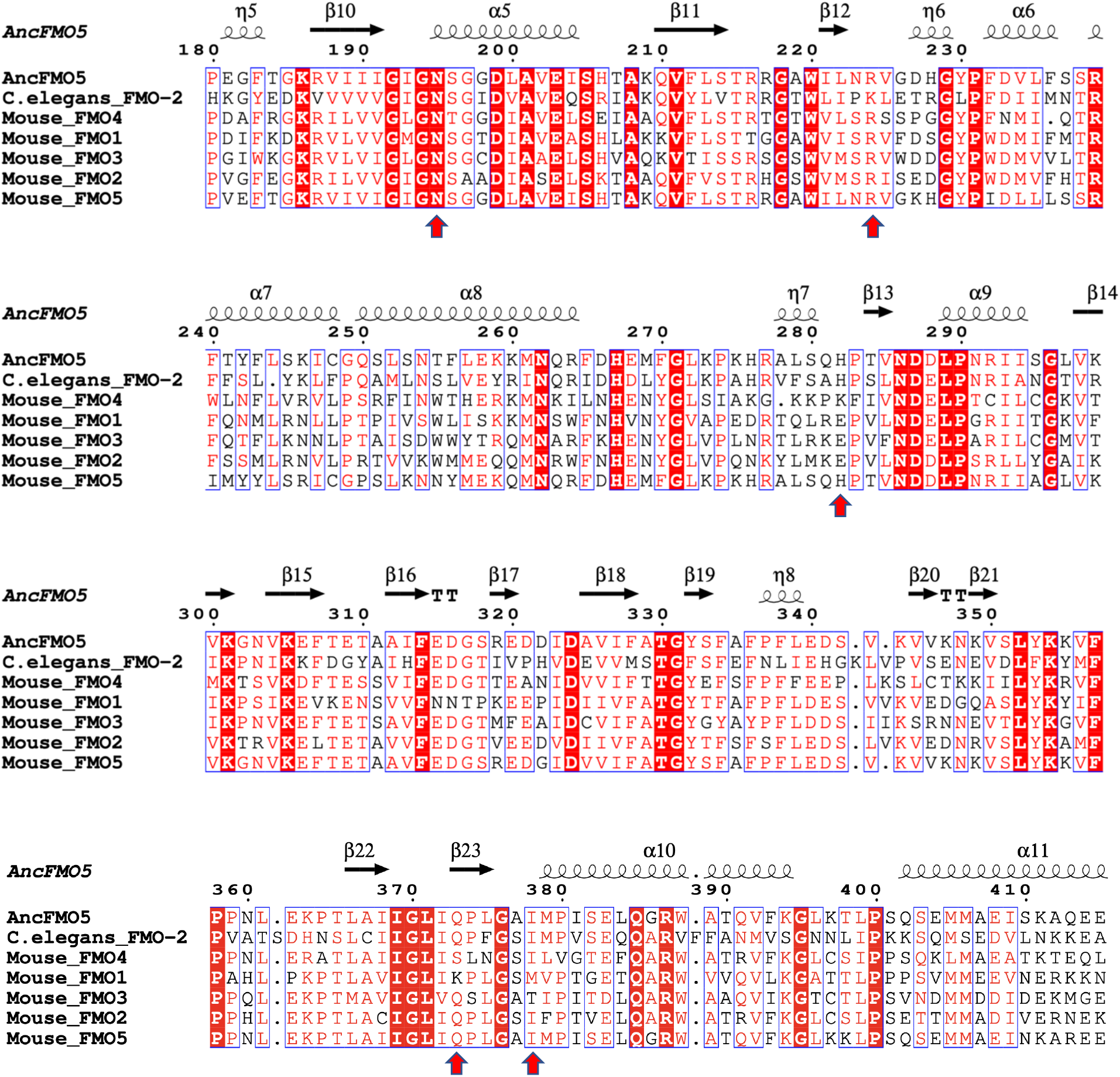
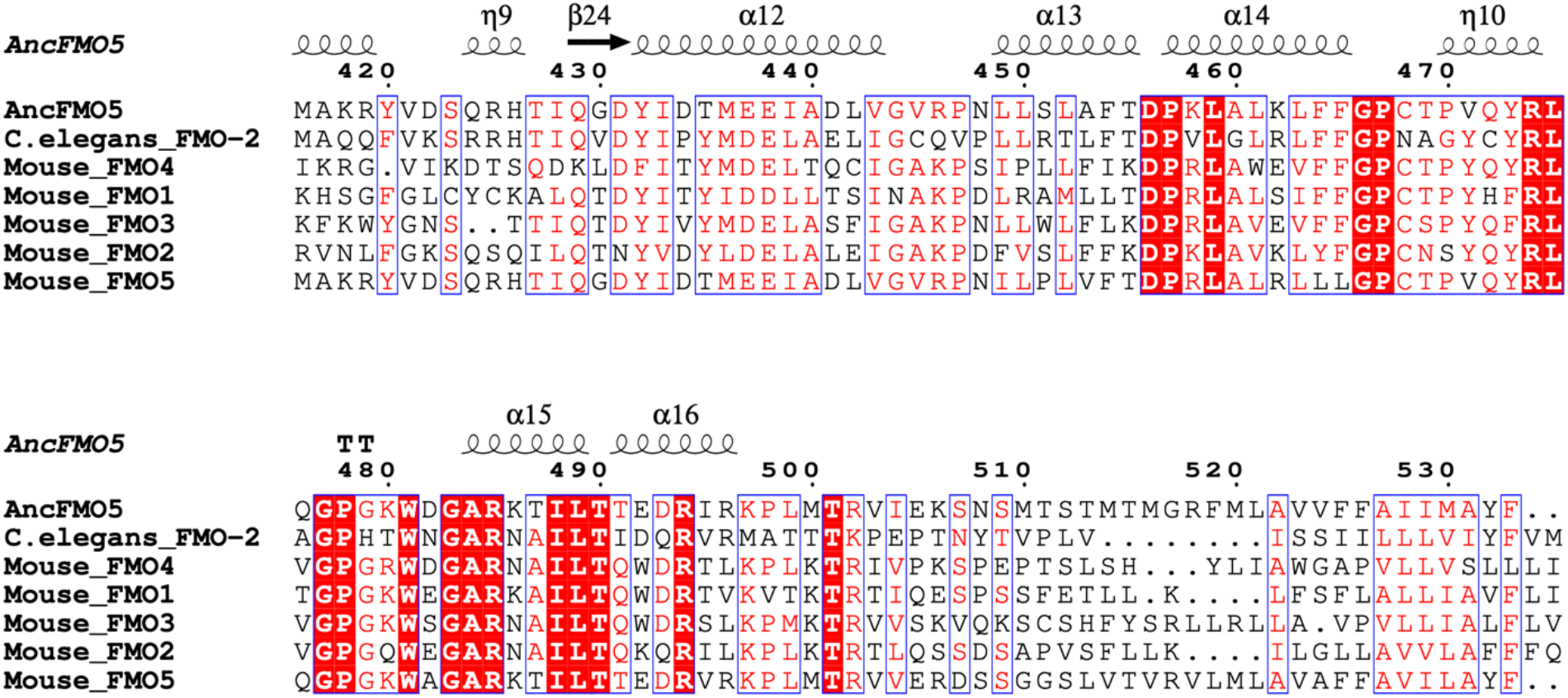
Conservation of catalytic residues between murine FMO5 and *C. elegans* FMO-2. Arrow heads indicate catalytic residues as determined by the crystal structure of ancestral mammalian FMO5 solved by Nicoll, et al. Overall identity of murine FMO5 and *C. elegans* FMO-2 is approximately 43%. Alignment figures were generated using Clustal Omega and ESPript 3.0.

To test this hypothesis, we assessed whether genes involved in tryptophan metabolism interact epistatically with FMO-2 (**Figure 6A**). Like our RNAi analyses of the OCM genes, we observed changes in stress resistance of the wild type and FMO-2 OE following the knockdown of genes involved in the kynurenine pathway (**Figures 6B, C**). Similarly, we observed changes in the lifespan of the wild type, FMO-2 OE, and FMO-2 KO worms under the same conditions (**Figures 6D, E**), as assessed using log-rank test with a cutoff threshold of p < 0.0001^28^ compared to the empty vector (EV) controls. Here, we again observed a separation between the regulation of stress resistance and lifespan under *kmo-1* and *tdo-2* knockdown (**Figures 6B-E**). Knocking down *kmo-1* increases the resistance to paraquat in the wild type (**Figure 6B**), but it decreases the lifespan of the wild type, FMO-2 OE, and FMO-2 KO (**Figure 6D**). These data suggest that *kmo-1* knockdown may be beneficial for resistance against paraquat, but that *kmo-1* expression is necessary for normal worm longevity. Knocking down *tdo-2* abrogates paraquat resistance in FMO-2 OE (**Figure 6C**), but extends the lifespan of the wild type, FMO-2 OE, and FMO-2 KO (**Figure 6E**). *Tdo-2* knockdown was previously reported to extend lifespan by inhibiting tryptophan degradation and thereby improving the regulation of proteotoxicity^19^.

**Figure 6:**
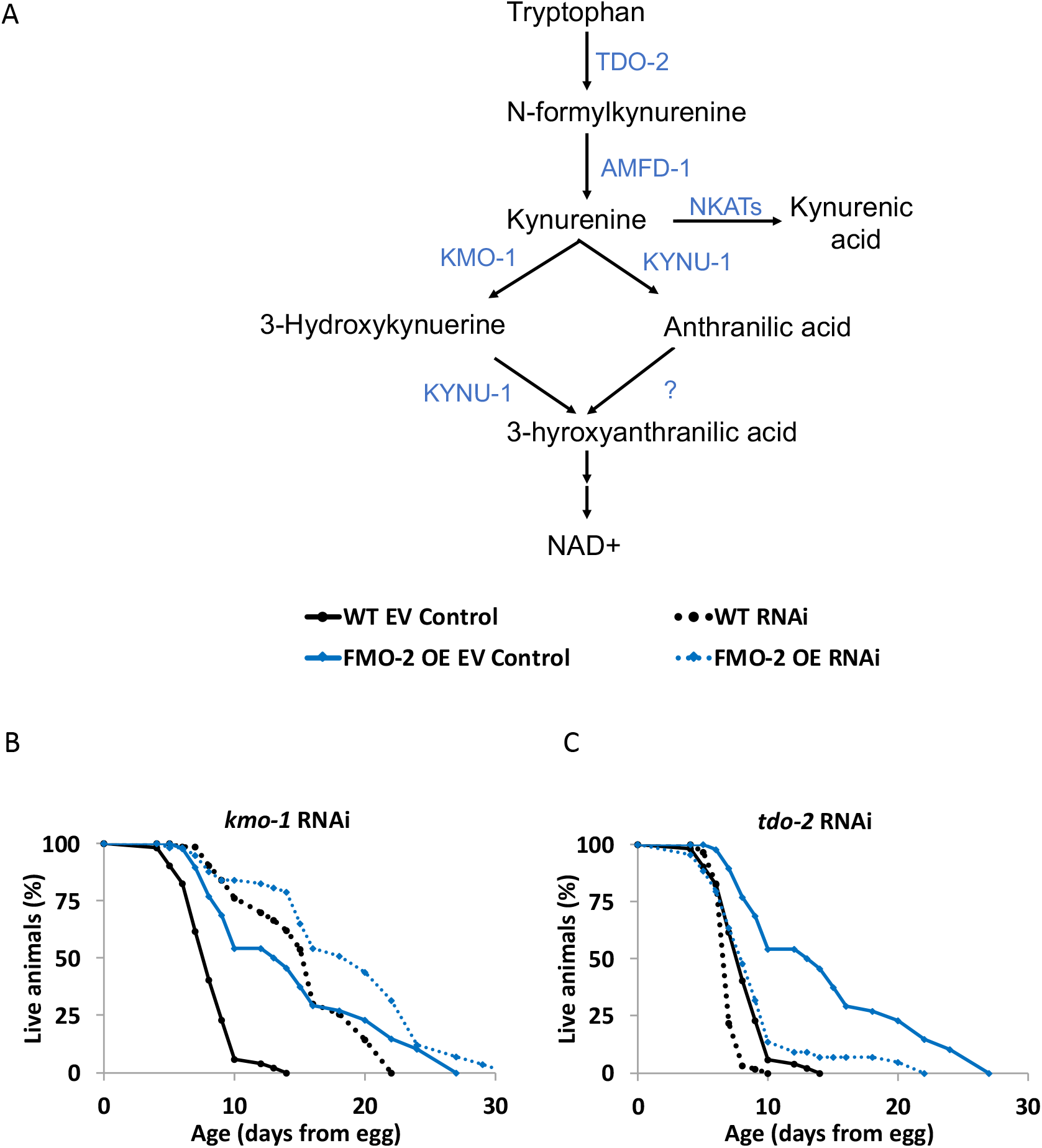
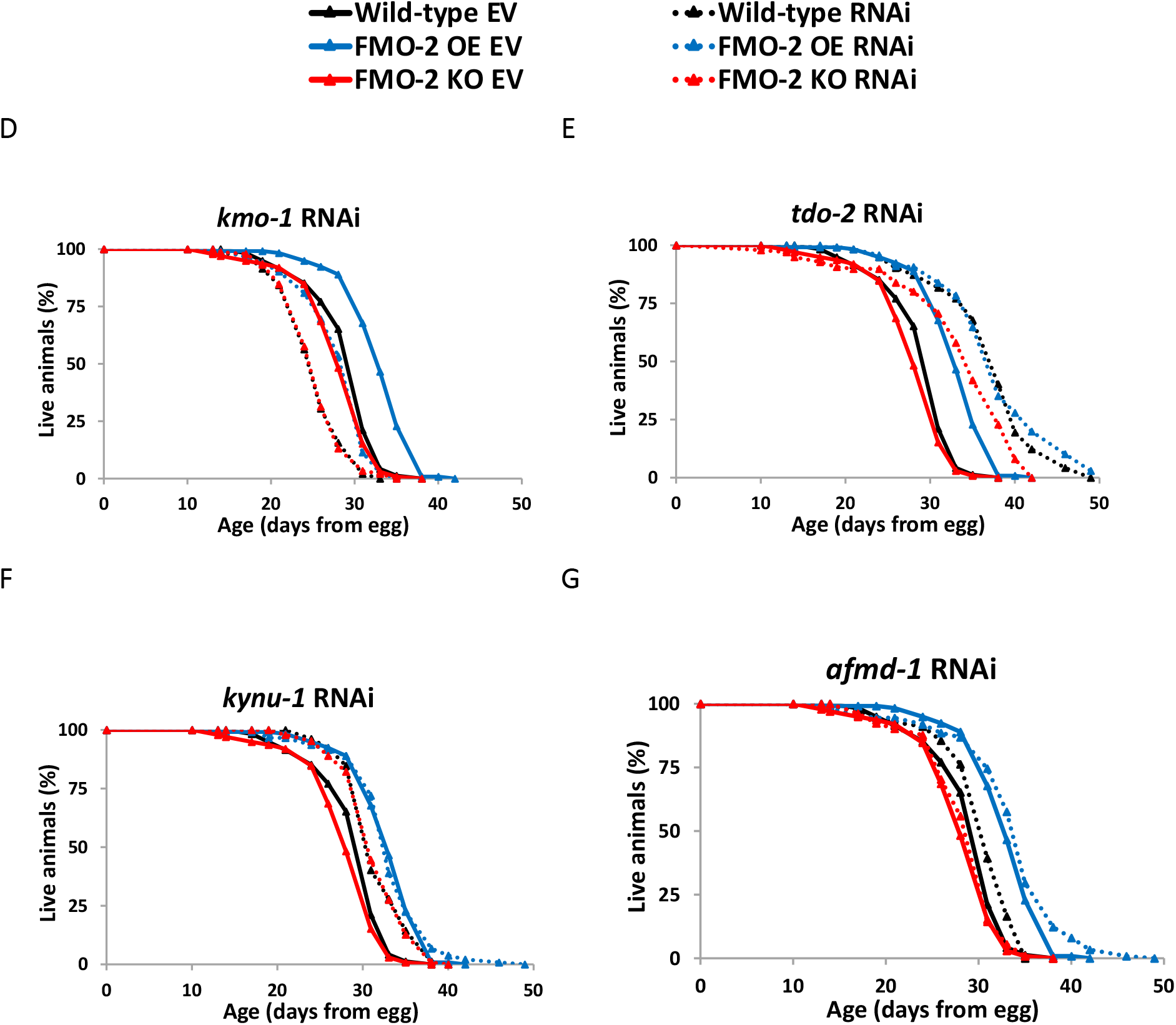
*Fmo-2* interacts with kynurenine metabolism to regulate stress resistance and lifespan. A) Diagram of kynurenine pathway. B-C) 5 mM paraquat stress assay (from L4 stage) comparing the survival of the wild-type and FMO-2 OE on empty vector (EV) and B) *kmo-1* RNAi and C) *tdo-2* RNAi, D-G) Lifespan assay comparing the survival of the wild-type, FMO-2 OE, and FMO-2 KO on EV and D) *kmo-1* RNAi, E) *tdo-2* RNAi, F) *kynu-1* RNAi, and G) *afmd-1* RNAi. Statistics in Supplementary Data 4 for stress assays and Supplementary Data 5 for lifespans.

In addition, knocking down *kynu-1* did not affect paraquat resistance of the worms (**Supplementary Figure 6A**), but it partially recapitulated the FMO-2 OE lifespan phenotype in the wild type and FMO-2 KO without affecting FMO-2 OE lifespan (**Figure 6F**), consistent with *kynu-1* functioning in the same pathway as *fmo-2*. The knockdown of *afmd-1* did not affect the stress resistance of the worms (**Supplementary Figure 6B**), but it extended the lifespan of the wild type without affecting FMO-2 KO and FMO-2 OE, suggesting that *afmd-1* requires *fmo-2* to extend lifespan (**Figure 6G**). Knockdown of *nkat-1* did not affect the lifespan or the paraquat resistance of the wild type, FMO-2 OE, and FMO-2 KO, suggesting that this gene does not function in the same pathway as *fmo-2* (**Supplementary Figures 6C, D**). Taken together, our data are consistent with the hypothesis that the kynurenine pathway is a target of FMO-2.

**Supplementary Figure 6:**
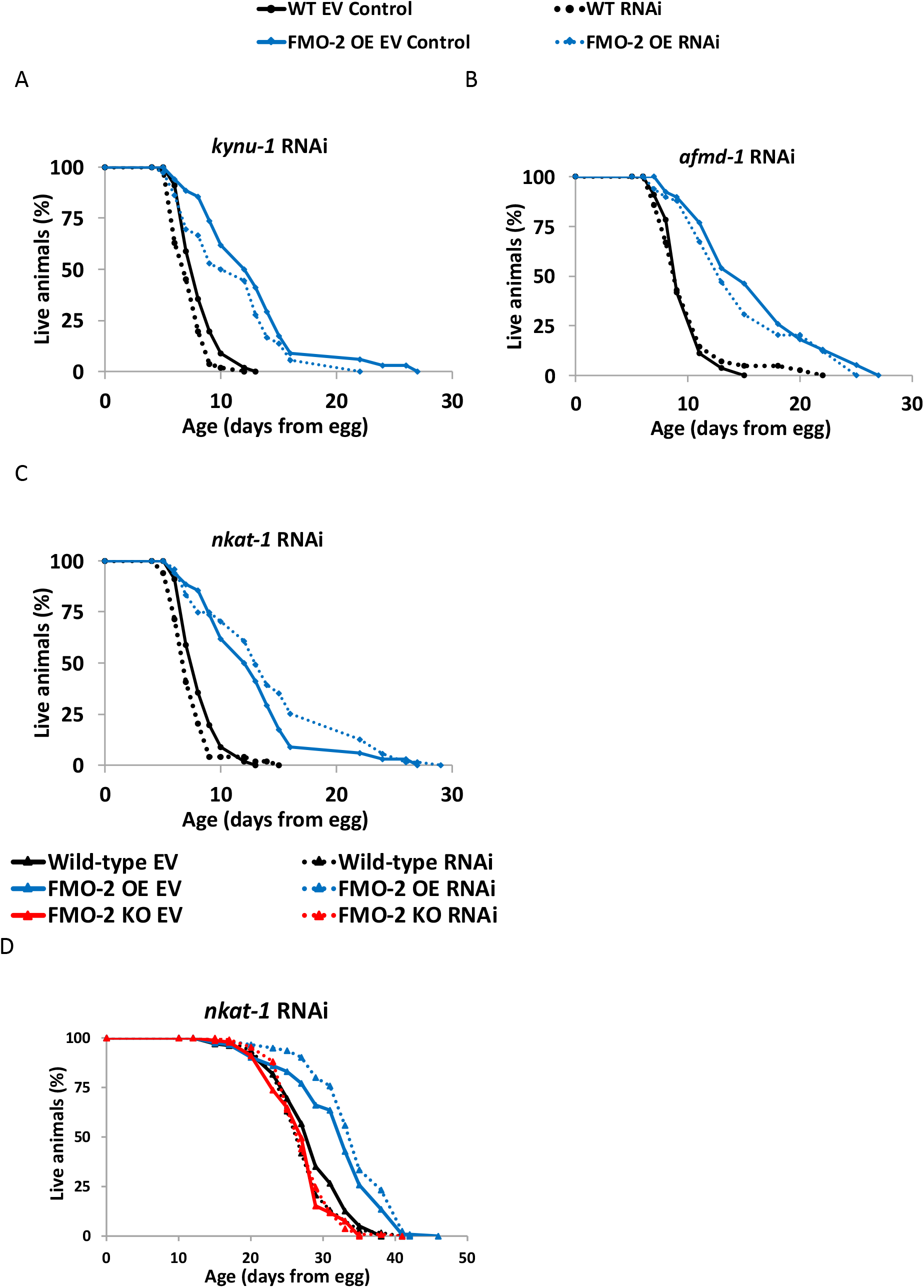
Kynurenine pathway genes that do not alter worm stress resistance or lifespan. A-C) 5 mM paraquat stress assay (from L4 stage) comparing the survival of the wild-type and FMO-2 OE on empty vector (EV) and A) *kynu1* RNAi, B) *afmd-1* RNAi, and C) *nkat-1* RNAi. D) Lifespan assay comparing the survival of the wildtype, FMO-2 OE, and FMO-2 KO on EV and *nkat-1* RNAi. Statistics in Supplementary Data 4 for stress assays and Supplementary Data 5 for lifespans.

## Discussion

For half a century, FMOs have been primarily classified as xenobiotic enzymes. However, the mechanisms by which these enzymes affect endogenous metabolism are still not well studied. Based on our data, we propose a model where overexpression of *fmo-2*, similar to levels that we observe under hypoxia and dietary restriction, is sufficient to remodel metabolism in the nematode *C. elegans* (**Figure 7**). Here, we show that *Cefmo-2*, a novel regulator of longevity that is critical for lifespan extension and stress response under dietary restriction and hypoxia, interacts with both tryptophan and one-carbon metabolism to confer longevity and health benefits. We find that modulating the expression of a single oxygenating protein can have a multitude of metabolic and physiological effects, similar to the activation of transcription factors and kinases. Our results suggest a broader, more significant role for FMO-2, and FMOs in general, than previously known. Furthermore, we establish experimental evidence of FMO orthology from *C. elegans* to mammals since both CeFMO-2 and mFMO5 have similar activity toward tryptophan, suggesting that through this substrate they may perform similar metabolic roles in both animals.

**Figure 7:**
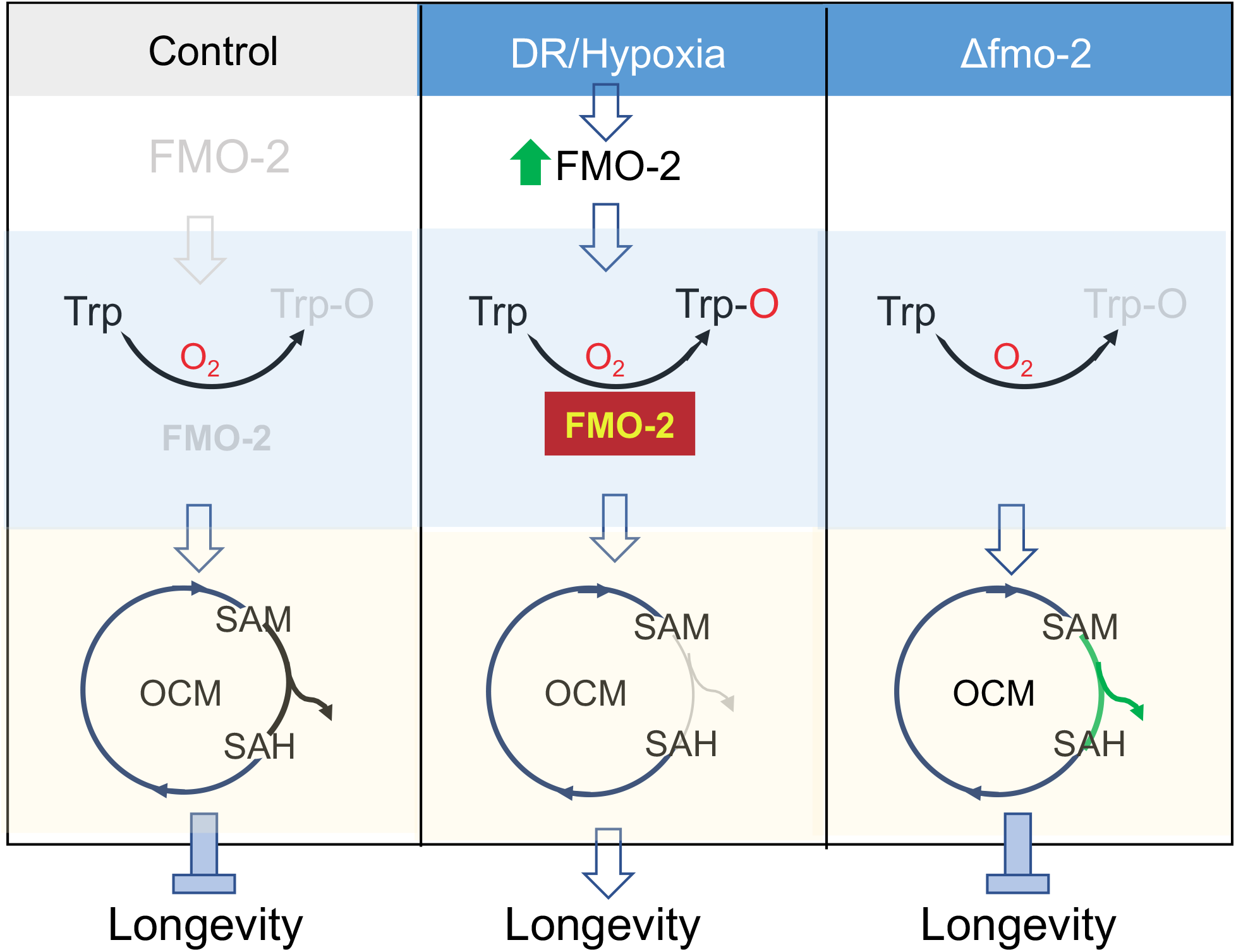
Proposed model. In control conditions, there is very low *fmo-2* expression, leading to low levels of tryptophan metabolism/kynurenine production through FMO-2, and maintaining normal flux through one carbon metabolism and normal lifespan. When *fmo-2* is induced, either genetically through overexpression or environmentally by DR or hypoxia, FMO-2 oxygenates tryptophan, leading to increased kynurenine production and decreased methylation output flux through OCM, thereby extending nematode lifespan. When *fmo-2* is absent, these metabolic changes do not occur, even under hypoxia or DR, preventing an extension in lifespan. The gray line represents decreased flux and the green line represents increased flux.

Our resulting data are consistent with a model where the reduction of flux through the methylation pathway leads to longevity benefits. By projecting gene expression data to a stoichiometric model for OCM metabolism, we predict that FMO overexpression results in a reduction in methylation flux. This model-based prediction based on gene expression data is experimentally validated, indicating that this approach can be a powerful tool to simplify the understanding of complex metabolic pathways and to study the biology of aging. Perturbation in the SAM/SAH ratio by either the supplementation of metformin or a mutation in *sams-1* extends worm lifespan^13,17^. While multiple studies report that methionine restriction robustly extends lifespan across species, including worms, flies, and mice^13,43,44^, others show that exogenous supplementation of methionine is not detrimental to lifespan^45^. This suggests that methionine utilization rather than methionine abundance is a key factor that influences the aging process.

Although suppressing *sams-1* expression phenocopies FMO-2 OE lifespan in the wild type and FMO-2 KO, doing so reduces the stress resistance of the worms against paraquat. This separation of lifespan and stress resistance is occasionally observed under other long-lived conditions^46^. It is unclear if simply reducing methylation is sufficient to promote longevity benefits, or if this mechanism requires suppression of specific methylation processes. It will be important for future studies to determine how cells regulate different methylation fluxes under *sams-1* knockdown and decreased overall methylation. One potential mechanism under this genetic condition could be that specific methyltransferases that are essential for survival will have higher affinity to methyl groups to outcompete other nonessential or deleterious methyltransferases.

We note that while our data suggest methylation as the key downstream effector of FMO-2, we have not excluded the possibility that the transsulfuration pathway may also be involved in this mechanism. The transsulfuration pathway is reported to be a necessary and sufficient component of DR-mediated lifespan extension in flies^16^. Similarly, knocking down *cth-2*, a gene involved in this pathway, abrogates the lifespan extension phenotype in FMO-2 OE (**Figure 3B**). It will be interesting to determine the mechanistic relationship between the transsulfuration and transmethylation pathways in regulating longevity.

Our data also support an interaction between *fmo-2* and tryptophan metabolism to influence longevity. These findings are particularly interesting because we identify a putative endogenous metabolic pathway of FMOs in relation to the aging process. Based on cell line metabolomics, enzyme kinetics, and HPLC data, there are at least two plausible mechanisms for how oxygenation of tryptophan by FMO-2 can lead to the synthesis of N-formylkynurenine. First, FMOs across taxa are known to dimerize and form higher order oligomers^47,48^. Therefore, it is possible that FMO-2 dimerizes and dioxygenates tryptophan forming N-formyl-kynurenine, which is then converted to kynurenine by formamidase. Second, the same process could be involved in subsequent oxygenation by FMO-2 monomers, but it is unknown how stable a monooxygenated form of tryptophan would be within the cell, making the first mechanism more likely. To our knowledge this is the first example of the dioxygenation of a substrate that could potentially require dimerized FMOs. The mechanism of this reaction and its potential requirement of dimerized FMOs will be a target of future research. Furthermore, the dioxygenation of tryptophan by FMOs is especially interesting considering only dioxygenases, such as *tdo-2*, *ido-1*, and *ido-2*, have been shown to mediate the conversion of tryptophan to N-formylkynurenine^19^. Regardless, our data implicate tryptophan as a *bona fide in vitro* and likely *in vivo* substrate of animal FMOs either through dioxygenating or monooxygenating mechanisms. Although we tested multiple conventional and unconventional FMO substrates, such as methimazole^47–49^ and 2-heptanone^50^ (**Supplementary Data 9**), respectively, much work remains to fully establish the general FMO-2 substrate profile and how it compares to those of mammalian FMOs beyond the common tryptophan substrate of CeFMO-2 and mFMO5.

This is the first report showing the conservation of enzymatic activity toward a potential endogenous substrate between *C. elegans* and mammalian FMOs. Although FMO5 has been posited as the ortholog of FMO-2 and the other *C. elegans* FMOs due to it being the most ancestral mammalian FMO, experimental evidence demonstrating this has been lacking. The alignment of *C. elegans* FMO-2 with mFMO5 illustrates the highly conserved putative catalytic residues amongst the FMOs (**Supplementary Figure 5**). This conserved activity is demonstrated further here in the similar activities of CeFMO-2 and mFMO5 toward tryptophan as well as the production of N-formylkynurenine from tryptophan. As we show here metabolomics analysis reveals that multiple human cell lines with overexpression of mFMOs have less tryptophan^10^. Altogether, our previous data and the data presented here further suggest that not only is tryptophan a *bona fide* endogenous target of C. elegans FMOs but potentially of mFMOs as well.

Our data support a model where the interaction between FMO-2 and tryptophan metabolism directly or indirectly modulates the metabolite profile of OCM, altering flux patterns that are consistent with our computational model predictions and subsequent genetic analyses. Further investigation is needed to understand the *fmo-2*-mediated connection between OCM and tryptophan in regulating lifespan. Based on the knowledge we gained from this study and previous work, we propose the following possibilities: 1) Oxygenation of tryptophan by FMO-2 alters OCM flux by increasing formate levels as a potential direct link between tryptophan metabolism and OCM. Formate is a single carbon-containing molecule that can enter the folate cycle as a carbon source^42^. Formate is generated as a byproduct when kynurenine is synthesized from N-formylkynurenine by formamidase^42^. It is possible that increased of formate levels can confer stress resistance and longevity benefits under metabolically stressful conditions, such as DR or hypoxia. 2) FMO-2 interacts with the mechanistic target of rapamycin (mTOR). Dietary restriction leads to inhibition of mTOR signaling, which is a central regulator of lifespan and aging^51^. Interestingly, both DR- and rapamycin-mediated mTOR inhibition induce the expression of FMOs. A recent study shows that diaminodiphenyl sulfone (DDS) induces the expression of *fmo-2* and extends lifespan, but it does not further extend lifespan in combination with rapamycin^37^. This finding is consistent with the hypothesis that *fmo-2* interacts with mTOR inhibition to extend lifespan. We show that *fmo-2* interacts with SAM and tryptophan metabolism, both of which are known to alter mTOR activity^52–54^. Thus, examination into the role of mTOR in *fmo-*2-mediated lifespan extension is warranted. 3) FMO-2 modulates tryptophan metabolism and OCM in an independent manner. There is a possibility that there is no connection between OCM and tryptophan, and FMO-2 targets both pathways independently.

Taken together, our study expands the role of FMO-2 from a xenobiotic enzyme to a metabolic regulator of longevity that has global effects on the metabolome in worms. In particular, the identification of OCM as a target of FMO-2 has implications outside the aging field, considering that OCM remodeling has been studied under the context of cancer biology for more than 70 years^55^. Furthermore, through the identification of tryptophan as a putative substrate for both CeFMO-2 and mFMO5, this study highlights the conserved importance of FMOs in multiple contexts, including aging and many diseases where OCM and/or the kynurenine pathway play a role. These findings illustrate the potential for therapeutic targets of these proteins for treating age-related diseases and/or increasing longevity and healthspan. This exciting translational potential for the conserved roles of FMOs will be a target for future research.

## Materials and Methods

### Strains and Growth Conditions

Standard *C. elegans* cultivation procedures were used as previously described^4,56^. N2 wild type, KAE9 ((*eft-3p*::*fmo-2* + *h2b*::*gfp* + Cbr-*unc-119*(+)), and VC1668 (*fmo-2(ok2147)*) strains were maintained on solid nematode growth media (NGM) using *E. coli* OP50 throughout life except where RNAi (*E. coli* HT115) were used. All experiments were conducted at 20°C.

### Metabolomics

OP50 bacteria were treated with 0.5% (v/v) paraformaldehyde as previously described^56^ and seeded onto 100 mm NGM plates. Approximately 500 eggs were put on these plates and grown until they reached late L4 larval stage. The worms were washed off the plates with 10 mL of M9 buffer and were collected in 15 mL conical tubes. The worms were pelleted using a clinical centrifuge for 1 minute at 150 x g and the supernatant was vacuum aspirated. The worms were washed once with 10 mL of M9 buffer and then with 10 mL of 150 mM ammonium acetate to remove phosphates from M9, each time being centrifuged and the supernatant being aspirated. After these washing steps, the pellets were flash frozen in liquid nitrogen.

Metabolites were extracted from pellets by addition of 500 μL of ice-cold 9:1 methanol: chloroform, followed immediately by probe sonication for 30 seconds with a Branson 450 Sonicator. The resulting homogenates were kept on ice for 5 minutes and were then centrifuged for 10 minutes at 4000 x g at 4°C. Supernatant was then transferred to autosampler vials for analysis. Hydrophilic interaction liquid chromatography-electrospray ionization mass spectrometry (HILIC-LC-ESI-MS) analysis was performed in negative ion mode using an Agilent 1200 LC system coupled to an Agilent 6220 time-of-flight mass spectrometer. Chromatography was performed as previously described^57,58^, with the exception that the Phenomenex Luna NH2 column used had dimensions of 150 mm x 1.0 mm ID, the flow rate was 0.07 mL/min, and the injection volume was 10 μL. Untargeted peak detection and alignment was performed using XCMS^59^.

The resulting metabolomics data were analyzed using Metaboanalyst 4.0 (http://metaboanalyst.ca). Within Metaboanalyst, the data were median normalized, adjusted using auto scaling, and were then subjected to principal component analysis using default parameters. Pathway analysis was performed using Metaboanalyst’s functional analysis module. P-values and t-scores of each MS peak data were calculated between the wild type and FMO-2 OE (**Supplementary Data 2**). Mass tolerance was set to 10 parts per million (ppm) and mummichog algorithm p-value cutoff was set to 0.05. Default parameters were used for other settings and the analysis was done using the *C. elegans* pathway library.

Targeted metabolomics analysis used the same LC-MS parameters as untargeted, but data analysis was performed using Agilent MassHunter Quantitative Analysis software. Metabolite identification was performed by matching accurate mass and retention time with authentic standards analyzed by the same method. Data were normalized to the median and log transformed. Statistical analysis for targeted metabolomics data was done using Metaboanalyst.

### Stress resistance assay

Paraquat (Methyl viologen dichloride hydrate from Sigma-Aldrich) was used to induce oxidative stress. Worms were synchronized from eggs on RNAi plates seeded with *E. coli* HT115 strain expressing dsRNAi for a particular gene and at L4 stage 40 worms were transferred on RNAi-FUDR plates containing 5 mM paraquat. A minimum of two plates per strain per condition were used per replicate experiment. Worms were then scored every day and considered dead when they did not move in response to prodding under a dissection microscope. Worms that crawled off the plate were not considered, but ruptured worms were noted and considered as previously described^4^.

### Lifespans

Gravid adults were placed on NGM plates containing 1mM β-D-isothiogalactopyranoside (IPTG), 25 μg/ml carbenicillin, and the corresponding RNAi clone from the Vidal or Ahringer RNAi library. After 3 hours, the adults were removed, and the eggs were allowed to develop at 20°C until they reached late L4/young adult stage. From here, 40 to 90 worms were placed on each RNAi plate and transferred to fresh RNAi + FUDR plates on day 1, day 2, day 4, and day 6 of adulthood. A minimum of two plates per strain per condition were used per replicate experiment. Experimental animals were scored every 2-3 days and considered dead when they did not move in response to prodding under a dissection microscope. Worms that crawled off the plate were not considered, but ruptured worms were considered as previously described^4^.

### Computational Modeling

The computer model was generated by building a stoichiometric matrix S (10 reactants by 13 reactions), accounting for all reactions shown in Fig 4A. A steady-state approximation was used, as shown in Eq 1. In Eq. 1, S is the stoichiometric matrix and **J** is a vector of fluxes for each of the reactions.

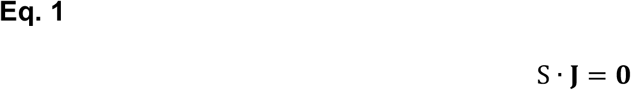

To obtain a biologically relevant solution, we projected the expression data of genes involved in the reactions used in the model to the nullspace of S by solving for Eq. 2. Single genes were used as representative genes for each reaction to simplify the model. Gene expressions related to input fluxes were assumed to be one for all strains. Reactions used in the model and the relevant gene expression data are shown in **Supplementary Data 7**. In Eq. 2, M is the nullspace of S, **b** is the vector of relative gene expression data from the wild type, FMO-2 OE or FMO-2 KO that have been normalized to the wild type, and **x** is a vector such that S**x** is the projection of **b** onto the column space of M, which gives us the vector of reaction fluxes, **J**, within the nullspace of S. To account for data variability, expression level with greater than 0.5x or less than 1.5x fold changes were assumed to be equal to the wild type control. Eq. 2 was solved using the lsqminnorm function in MATLAB 2018a. The lsqminnorm function returns the minimum norm least-squares solution to M**x** = **b** by minimizing both the norm of M * **x** – **b** and the norm of **x**.

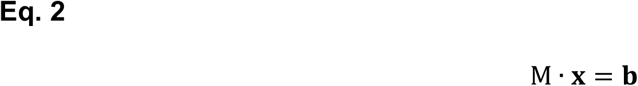

The inner product of the resulting vector **x** and the nullspace matrix M was obtained to calculate the reaction flux predictions resulting from the gene expression projection as shown in Eq. 3. The calculated **J** for FMO-2 OE and FMO-2 KO were normalized to that of the wild type to obtain the relative fluxes.

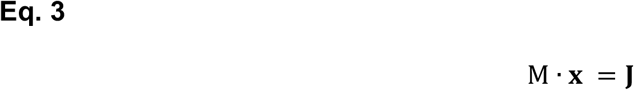

### Quantitative PCR

RNA was isolated from day 1 adult worms following three rounds of freeze-thaw in liquid nitrogen using Invitrogen’s Trizol extraction method and 1 μg of RNA was reverse transcribed to cDNA using SuperScript^™^ II Reverse Transcriptase (Invitrogen). Gene expression levels were measured using 1 μg of cDNA and SYBR^™^ Green PCR Mastermix (Applied Biosystems) and primers at 10 μM concentration. mRNA levels were normalized using previously published housekeeping gene controls, *tba-1* and *pmp-3*^60^. List of primers used are in **Supplementary Data 11**.

### Enzyme Kinetic Assays

Oxygenation activity of FMO-2 and mFMO5 was characterized using the method previously described^61^. Briefly, oxygenation of substrates was determined by spectrophotometrically following the consumption of NADPH at 340 nm using the molar extinction coefficient 6.22 mM^-1^cm^-1^. Components of the assay buffer included 25 mM sodium phosphate buffer (pH 8.5), 0.5 mM diethylenetriaminepentaacetic acid (DETAPAC), 0.5 mM NADPH, and 0.04 μM FMO-2 (0.4 μM FMO5) with excess FAD. The final substrate concentrations for tryptophan were 100, 250, 500, 750 μM and 1, 2.5, 5, 7.5, and 10 mM. The final substrate concentrations for MMI were 100, 300, and 600 μM and 1, 3, 5, 7, 10, and 30 mM. To determine the rate of oxidation of NADPH by FMO, NADPH concentrations of 10, 30, 100, 300, 500, and 700 μM and 1 and 1.5 mM were used. Experiments were conducted at 30°C while shaking. Kinetic parameters (i.e *k_cat_* and *K_m_*) were determined by fitting plots of the rate of turnover vs the substrate concentration to the Michaelis-Menton equation using GraphPad Prism (version 9.1.0; GraphPad Software Inc., San Diego, CA.). Purified FMO-2 protein was purchased from GenScript. Purified FMO5 protein, NADPH, FAD, MMI, L-tryptophan, and all other substrates were purchased from Sigma Aldrich (St. Louis, MO). DETAPAC and sodium phosphate buffer were purchased from Fisher (Waltham, MA).

### *In vitro* studies LC-MS

Analysis of samples from *in vitro* studies with purified FMO2 and FMO5 protein was performed using LC-MS with untargeted feature detection. Samples contained 100, 250, or 500 μM tryptophan in the same conditions as the enzymatic assays with either FMO-2 or FMO5 proteins. 100 μL of conditioned media were vortexed with 400 μL of 1:1:1 methanol:acetonitrile:acetone to precipitate protein. The extract was centrifuged for 10 minutes at 16,000 x g and 200 μL of supernatant were transferred to a clean autosampler vial with insert and dried under a stream of nitrogen gas. The dried extract was reconstituted in 50 μL of 85/15 acetonitrile/water and analyzed by HILIC-TOF-MS on an Agilent 1290 Infinity II / Agilent 6545 QTOF. Chromatography was performed on a Waters BEH Amide column (2.1 mm ID x 10 cm, 1.7 μm particle diameter) with mobile phase prepared as described previously^62^ except that mobile phase A contained 5% acetonitrile. The flow rate was 0.3mL/min, the column temperature 55 °C, and the gradient was as follows: 0-0.70 min 100%B, 0.7-6.7 min 100-85%B, 6.7-8.7 min 85%B, 8.7-16 min 85-28%B, 16-16.7 min 28%B, 16.7-16.8 28-0%B. Total run time was 22 min. Ion polarity was positive, gas temp was 320 °C, drying gas was 8L/min, nebulizer was 35 psi, sheath gas temp and flow were 350 °C and 11 L/min, capillary voltage 3500V. The instrument was operated in full scan mode at 2 spectra/sec and a mass range of 50-1200 Da. Feature detection and alignment was performed using XCMS. Potential reaction products were detected by computationally examining the data for features present in each sample set. Identification of potential reaction products was performed using MS/MS data acquired from a pooled sample.

### Statistical analyses

Log-rank test was used to derive p-value for lifespan and paraquat survival assays using p < 0.0001 cut-off threshold compared to EV controls. Paired-test was used to derive p-values for targeted metabolomics data using p < 0.05 cut-off threshold compared to the wild type. One-way ANOVA followed by Tukey’s *post hoc* test was used to derive p-values for SAM/SAH ratio using p < 0.05 cut-off threshold. Paired t-test was used to derive p-values for comparing the metabolomics data of HepG2 pDEST control and FMO2, FMO4, and FMO5 OE cell lines using p < 0.05 cut-off threshold.

