## Supplementary material for "FMO rewires metabolism to promote longevity through tryptophan and one carbon metabolism": Supplementary Data 6.docx

```
% Please replace the spaces in Supplementary Data file names with underscores to be able to use
% the script.

S = xlsread('Supplementary_Data_8_Stoichiometric_matrix.xlsx','B2:N10');  %Reads one carbon metabolism stoichiometric matrix

M = null(S,'r'); %Nullspace of S

fc_ox = xlsread('Supplementary_Data_7_model_reactions_and_gene_expression.xlsx','D2:D14'); %Reads fmo-2 overexpression relative gene expression fold change
fc_ko = xlsread('Supplementary_Data_7_model_reactions_and_gene_expression.xlsx','E2:E14'); %Reads fmo-2 knockout relative gene expression fold change
fc_wt = ones(13,1);

for i = (1:length(fc_ox)) %Cut-off for gene expression data
if fc_ox(i) < 1.5 && fc_ox(i) > 0.5
   fc_ox(i) = 1;
end
end

for i = (1:length(fc_ko)) %Cut-off for gene expression data
if fc_ko(i) < 1.5 && fc_ko(i) > 0.5
   fc_ko(i) = 1;
end
end

a_ox = lsqminnorm(M,fc_ox)'; %relative factor for fmo-2 overexpression strain
a_ko = lsqminnorm(M,fc_ko)'; %relative factor for fmo-2 knockout strain
a_wt = lsqminnorm(M,fc_wt)'; %relative factor for wild type

J_ox = M * a_ox'; %flux prediction for fmo-2 overexpression strain
J_ko = M * a_ko'; %flux prediction for fmo-2 knockout strain
```

```
J_wt = M * a_wt'; %flux prediction for wild type

J_rel_ox = J_ox./J_wt; %relative flux prediction for fmo-2 overexpression strain
J_rel_ko = J_ko./J_wt; %relative flux prediction for fmo-2 knockout strain


figure
cdata = [J_rel_ox,J_rel_ko];
xvalues = {'FMO-2 OE','FMO-2 KO'};
yvalues = {'met -> sam','sam -> sah','sah -> hcy','hcy + 5mthf -> met + thf',...
    'hcy -> cyst','cyst -> cys','thf -> 5,10thf','5,10thf -> 5mthf','METin',...
    'FOLin', 'methylation','cys consumption','pyrimidine synthesis'};
h = heatmap(xvalues,yvalues,cdata);
```
